# Multi-omics and biophysical phosphoproteomics upon BRAF inhibition uncover functional networks of BRAFV600E-driven signaling

**DOI:** 10.64898/2026.02.09.704793

**Authors:** Mira Lea Burtscher, Martin Garrido-Rodriguez, Pablo Andres Rivera Mejias, Dimitrios Papagiannidis, Isabelle Becher, Denise Medeiros Selegato, Clement Potel, Ferris Jung, Michael Zimmermann, Julio Saez-Rodriguez, Mikhail M Savitski

**Affiliations:** European Molecular Biology Laboratory (EMBL), Heidelberg, Germany; Faculty of Biosciences, Heidelberg University, Heidelberg, Germany; Heidelberg University, Faculty of Medicine, and Heidelberg University Hospital, Institute for Computational Biomedicine, Heidelberg 69120, Germany; European Molecular Biology Laboratory, European Bioinformatics Institute (EMBL-EBI), Hinxton CB10 1SD, United Kingdom

## Abstract

Dysregulated kinase activity drives oncogenic signaling, disrupts cellular homeostasis, and promotes tumour progression. The BRAFV600E mutation constitutively activates the MAPK pathway and is a key therapeutic target in melanoma and other cancers, but the functional relevance of most downstream phosphorylation events and mechanisms of drug resistance remain unclear. To address this, a global multi-omic model of BRAF inhibition response was established in BRAFV600E-mutant cells by integrating time-resolved and biophysical phosphoproteomics, transcriptomics, and thermal proteome profiling. Ultradeep phosphoproteomics revealed extensive phosphorylation changes upon BRAF inhibitor treatment, while biophysical phosphoproteomics identified phosphorylation events linked to altered protein solubility and subcellular localization, suggesting changes in nucleic acid interactions and nuclear reorganisation. Network-based integration of these datasets prioritized functionally relevant phosphorylation sites and kinases. Experimental validation identified CDK9, CLK3, and TNIK as critical regulators of BRAFV600E signaling and candidate targets for combinatorial inhibition capable of re-sensitising resistant cells. The transcription factor ETV3 emerged as a previously unrecognised effector of BRAF signaling. Biophysical proteomics data confirmed that ETV3 phosphorylation modulates DNA-binding, while functional assays combining knockdown, metabolomics, and drug screening demonstrated its role in coordinating transcriptional and metabolic adaptations to BRAF inhibition. This study provides a systems-level framework linking phosphorylation dynamics to protein function and phenotype, identifies ETV3 as a new node in oncogenic BRAF signaling, and illustrates how integrated, site-resolved models can reveal mechanisms of kinase-driven oncogenesis.

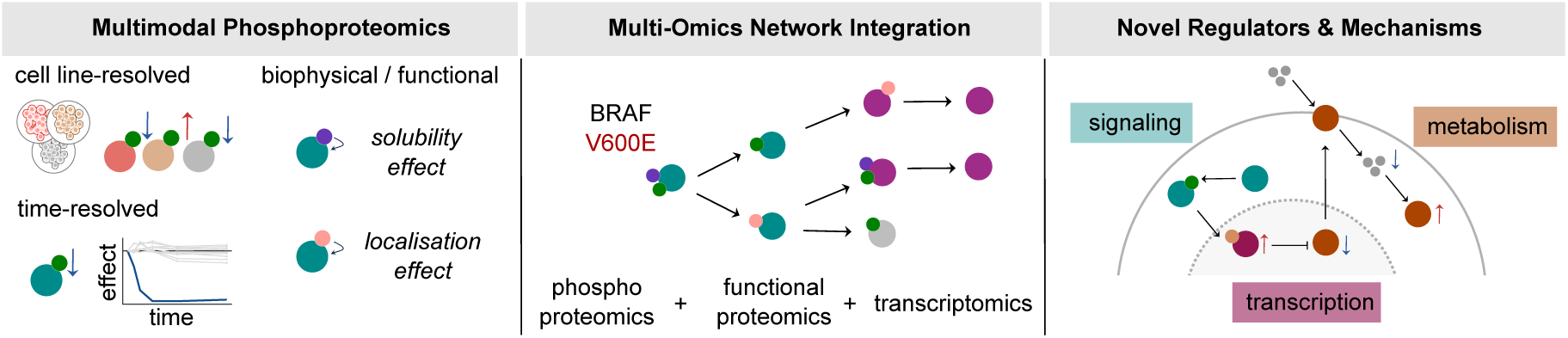

**Highlights:**

- Time- and cell type-resolved phosphoproteomics maps BRAF inhibition dynamics
- Biophysical phosphoproteomics, combining quantitative phosphoproteomics with solubility profiling or nuclear fractionation, reveals phosphorylation-driven changes of protein solubility and localization
- Integration of abundance and biophysical phosphoproteomics data identifies functionally relevant phosphorylation events of BRAFV600E signaling
- Network integration of multimodal phosphoproteomic, transcriptomic and thermal proteome profiling data links signaling to protein function and cellular phenotypes
- Biophysical evidence improves models and identifies non-canonical kinases driving BRAF signaling as well as novel downstream regulators such as ETV3
- Follow-up experiments reveal a ETV3-GLUT3-mediated metabolic adaptation in BRAFV600E cells

## Introduction

Reversible protein phosphorylation is one of the most pivotal and extensively studied post-translational modifications (PTMs) in biological systems, playing a central role in virtually all biological processes ^1^. Phosphorylation holds particular significance in oncology and cancer research for several reasons: (i) it governs critical signaling pathways that drive hallmark cancer behaviors such as uncontrolled proliferation, invasion, and metastasis ^2^; (ii) enzymes that catalyze phosphorylation, kinases are key therapeutic targets in cancer treatment^3^; and (iii) the dynamic reprogramming of signaling networks affects drug responsiveness and underlies the emergence of resistance mechanisms ^4^.

Hyperactive kinases driving oncogenic signaling are a key example of how deregulated phosphorylation perturbs cellular phenotypes and drives cancer development. Although many highly effective kinase inhibitors have been developed to specifically target these oncogenic signaling processes ^5^, approximately 30% of current drug development efforts continue to focus on this area ^6^. The B-Rapidly Accelerated Fibrosarcoma Kinase *V600E* mutation (BRAFV600E), prevalent in up to 70% of melanomas and recurrent in other cancers such as colorectal carcinoma, represents a prominent example of kinase hyperactivation in cancer^7^. This point mutation affects a key autoactivating phosphosite in the BRAF kinase, leading to its constitutive activation ^8^. BRAF functions upstream of Mitogen activated protein kinase (MAPK) signaling, and its deregulation drives altered cell proliferation ^9^. Several inhibitors, such as Dabrafenib ^10^, targeting this hyperactive form of BRAF have been developed and successfully implemented in cancer therapy. However, the detailed downstream signaling consequences as well as cytotoxic and resistance mechanisms of these inhibitors remain incompletely understood ^11–13^.

A major barrier for understanding oncogenic signaling is the limited functional annotation of phosphorylation events and their corresponding enzymes. Currently, upstream kinases have been identified for fewer than 5% of known human phosphorylation sites, and less than 3% of these sites have an assigned biological function ^14^. Therefore, comprehensive datasets that can provide systematic insights into the functional relevance of signaling alterations induced by mutant BRAF activity would be of great benefit for refining therapeutic strategies ^15^.

Here, we established a global model of the BRAF inhibition response in BRAFV600E-mutant melanoma cells by combining ultradeep phosphoproteomics with biophysical PTM-proteomics, transcriptomics and thermal proteome profiling (TPP) data. Biophysical proteomics approaches such as thermal ^16^ and solubility proteome profiling ^17^ (SPP) can provide mechanistic insight into how phosphorylation influences protein conformation, interaction, and localization, capturing potential functional effects beyond protein abundance changes ^18,19^. We generated biophysical phosphoproteomics datasets and identified phosphorylation events potentially involved in biomolecular condensation and subcellular relocalization. Integrating all modalities into a multi-omics network model enabled the formulation of mechanistic hypotheses linking altered phosphorylation dynamics to downstream signaling and functional outcomes at the phosphosite level.

Our multi-omics data, combined with biophysically weighted network modelling, revealed CDK9, CLK3, and TNIK as kinases that influence sensitivity to BRAF inhibition in a synergistic or synthetically lethal manner. Further, we identified the transcription factor ETS variant transcription factor 3 (ETV3) as a previously unrecognized regulator of melanoma cell response to targeted therapy and delineated a central role for ETV3 in mediating metabolic rewiring downstream of BRAFV600E signaling.

## Results

### Ultradeep phosphoproteomics reveals key axes of BRAFV600E-mutant driven signaling

To advance our understanding of oncogenic signaling mechanisms in BRAFV600E-driven cancers, we investigated three cell lines carrying this mutation (dataset overview in Supplementary Table 1). The experiments included two melanoma cell lines: A2058, which exhibits resistance to BRAF inhibition ^20^ (100 nM Dabrafenib around estimated plasma concentration of 190-960 nM ^10^) and MNT-1, which is sensitive to BRAF inhibition, as well as the colorectal cancer line COLO-201, which also shows sensitivity to BRAF inhibition. The three cell lines were treated with Dabrafenib (100 nM, 4 h) followed by quantitative phosphoproteomic profiling to capture the immediate signaling response (Figure 1A, Supplementary Table 2). Principal components analysis (PCA) revealed a mostly cell-line-specific response, but also a global, potentially context-independent signaling signature (Figure S1A). For instance, phosphorylation of the major oncogenic transcription factor MYC at S62, a known activating site ^21^, was found to be downregulated in all three BRAFV600E-mutant cell lines (Figure S1B). In general, phosphorylation signaling was downregulated in a global manner in response to Dabrafenib across all cell lines (Figure 1B), while protein abundance remained largely unchanged (Figure S1C), underscoring the central role of phosphorylation during this early response. Globally affected phosphoproteins showed a coherent signal across all cell lines for known BRAF regulated processes (Figure 1C). Nonetheless, 85% of phosphorylation events that were significantly altered by treatment occur in only one cell line suggesting strong context-dependency of BRAF inhibition (Figure 1D). To resolve the temporal signaling mechanisms underlying the resistance phenotype, we performed an ultradeep phosphoproteomic time course in A2058 cells (the least Dabrafenib-sensitive BRAFV600E model in our panel) and monitored phosphorylation dynamics of 85,109 phosphopeptides over time (Figure 1E, Supplementary Table 2). This dataset allowed us to track MAPK1 Y187 and MAPK3 Y204 phosphorylation and thus their inactivation over time, key events of the BRAF-MAPK signaling cascade (Figure S1D). Globally, the temporal response was dominated by a profound reduction of phosphorylation, with 1,192 phosphopeptides significantly downregulated and only 364 significantly upregulated after 240 minutes (moderated t-test, adj. p-value < 0.05, |log₂ fold-change| > log₂(1.5); Figure 1F, Figure S1E). To further dissect BRAFV600E-mutant signaling dynamics, we performed neural gas clustering ^22^ on phosphopeptides showing significant regulation in at least one time point (Figure 1G). This identified several distinct clusters of phosphorylation profiles, each associated with specific biological processes based on pathway enrichment analysis (p-value < 0.05, normalized weighted mean test; Figure 1H): (i) Cluster 1: Moderate downregulation of phosphorylation on proteins involved in RNA processing and splicing; (ii) Cluster 2: Strong downregulation on canonical MAPK signaling targets, including proteins involved in cell cycle regulation; (iii) Cluster 3: Moderate upregulation of phosphosites on cytoskeletal proteins, likely reflecting cytoskeletal disassembly processes; (iv) Cluster 4: Early and sharp phosphorylation (at 15 minutes) of proteins involved in heat shock response and chromatin remodeling. Closer inspection of the clusters revealed robust phosphorylation changes in key oncogenic regulators over time (Figure 1I). Overall, the obtained phosphoproteomic data reveal a complex landscape of BRAF inhibitor-induced signaling, with common regulatory events but distinct temporal patterns and high cell line specificity that likely contribute differences in drug sensitivity between BRAFV600E-mutant cell types.

**Figure 1:**
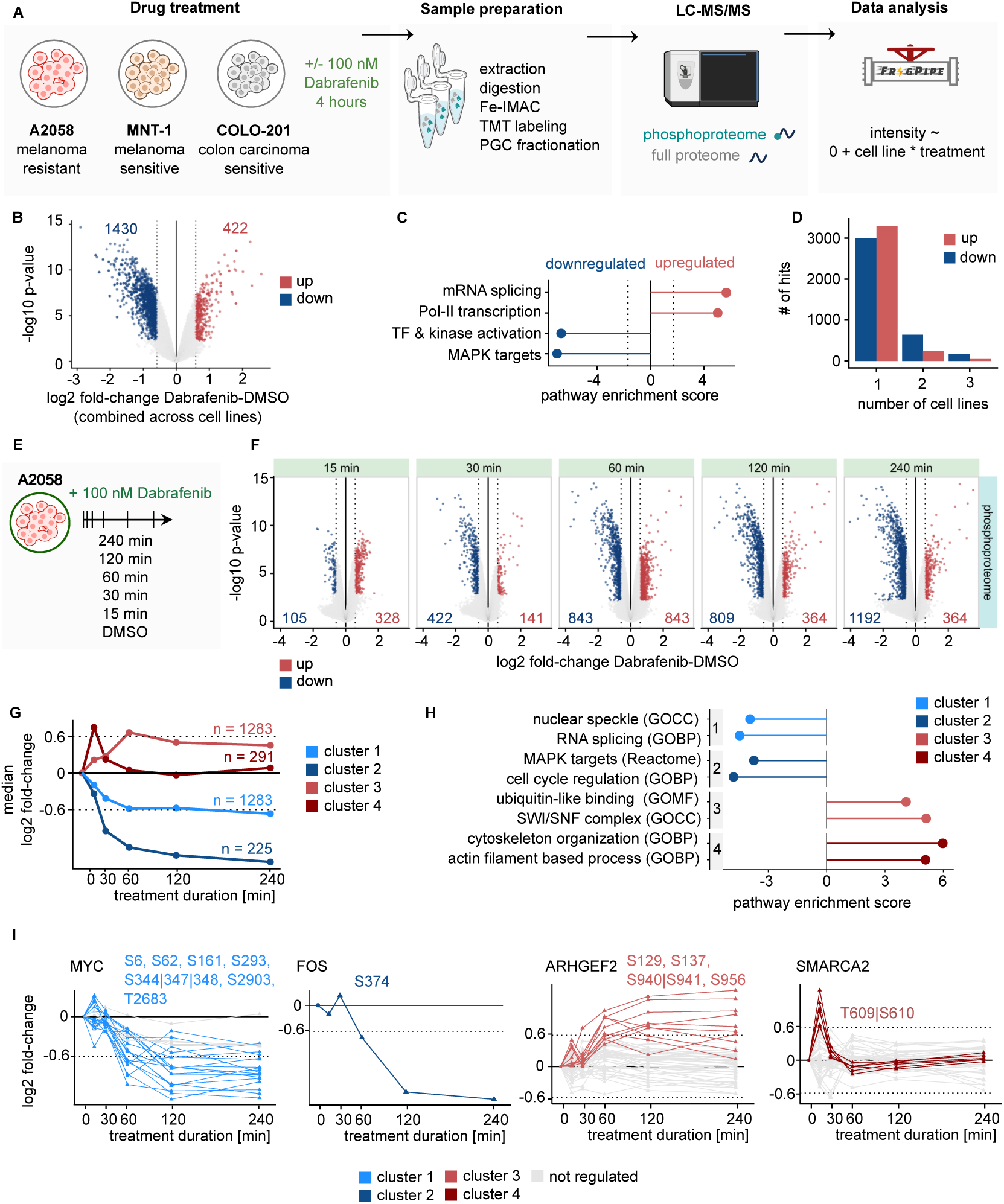
Ultradeep phosphoproteomics reveals key axes of BRAFV600E-mutant driven signaling. **(A)** Experimental workflow for Dabrafenib treatment (100 nM, 4 hours) in BRAFV600E-mutant cell lines (A2058, COLO-201, MNT1), followed by phosphoproteomic and proteomic profiling. Phosphopeptides were enriched using an automated Fe³⁺-IMAC platform and quantified by TMT18-plex labeling. Both phosphoproteome and full proteome samples were prefractionated using porous graphitized carbon (PGC) prior to LC-MS/MS analysis. Default data processing included database search with fragpipe, variance-stabilizing normalization and differential expression analysis with limma. **(B)** Volcano plots showing differential phosphopeptide abundance upon Dabrafenib treatment (100 nM, 4 h) relative to DMSO control across all three cell lines combined (A2058, COLO-201, MNT1). Significantly upregulated (red) and downregulated (blue) phosphopeptides are highlighted (moderated t-test, adj. p-value < 0.05, |log₂ fold-change| > log₂(1.5) as indicated by dashed line). **(C)** Pathway enrichment analysis of genes with up- or downregulated phosphopeptides upon Dabrafenib treatment across cell lines (normalized weighted mean test, Reactome pathways, p-value < 0.05, selected top pathways). **(D)** Number of phosphopeptide hits, stratified by direction of change and by the number of cell lines in which they are significantly altered in abundance upon Dabrafenib treatment (100 nM, 4h). **(E)** Experimental design of the Dabrafenib treatment time course experiment. A2058 melanoma cells were treated with 100 nM Dabrafenib or DMSO control and harvested at six time points (0, 15, 30, 60, 120, 240 min) in three biological replicates in reverse order. **(F)** Volcano plots showing differential phosphopeptide abundance upon Dabrafenib treatment (100 nM) at each time point relative to DMSO control at 0 hours. Significantly upregulated (red) and downregulated (blue) phosphopeptides are highlighted (moderated t-test, adj. p-value < 0.05, |log₂ fold-change| > log₂(1.5) as indicated by dashed line). **(G)** Temporal dynamics of four phosphopeptide clusters identified by neural gas clustering based on phosphopeptides significantly affected by Dabrafenib treatment in at least one time point. Line plots show median log₂ fold changes over time per cluster. Cluster sizes are indicated. **(H)** Pathway enrichment analysis enrichment analysis for genes of each phosphopeptide cluster (normalized weighted mean test with GO terms and Reactome pathways, p-value < 0.05, selected top pathways). **(I)** Representative temporal phosphorylation profiles of regulatory proteins of the different clusters, including MYC (multiple sites), FOS (S374), ARHGEF2 (S129, S137, S940/S941, S956), and SMARCA4 (T609/S610).

### Biophysical phosphoproteomics to investigate aspects of phosphorylation-driven protein regulation

While the insights gained from temporal and cell type-specific signaling analyses provide a valuable perspective on BRAF inhibition, identifying key phosphorylation events from thousands of affected phosphosites that determine cellular phenotypes remains a major challenge. To move beyond established functional annotations, we employed biophysical and nuclear enrichment phosphoproteomics to systematically assess the impact of phosphorylation on protein function. Solubility is a biophysical property that can inform on protein condensation state and nucleic acid interactions ^17^. To investigate which phosphorylation sites impact solubility, we performed an in depth phospho-SPP ^18^ in A2058 cells (solubility proteome profiling at the phosphopeptide level; Figure 2A, Supplementary Table 2). We profiled the solubility of 52,716 phosphopeptides and their corresponding proteins. On average, protein solubility was ∼78%, with no major difference observed between phospho- and non-phosphoproteins (Figure S2A). Consistent with pathway annotations and predictive models of protein condensation ^23^, proteins with low solubility, especially phosphoproteins, were significantly enriched in predicted condensate-forming features (p-value < 0.05, two-sided, unpaired t-test; Figure S2B, C). By comparing the solubility of a protein (reflecting the average protein population) with that of corresponding phosphopeptides, we were able to identify phosphorylation events that may regulate protein solubility and thus functions such as condensation and nucleic acid interaction. For instance, TOP2A, a topoisomerase with relatively low solubility, showed distinct behavior across phosphosites (Figure 2B). Phosphorylation at S1504 did not alter protein solubility, whereas phosphorylation at S1106 substantially decreased solubility, consistent with its known role in stabilizing DNA-protein complexes and promoting enzymatic activity ^24,25^. Another example is ARHGEF10, a RHO guanine nucleotide exchange factor potentially important in cytoskeletal regulation and signaling ^26^. While phosphorylation at S198 and S200 lacks prior functional annotation, our data indicate that only the doubly phosphorylated form affects protein solubility, suggesting potential involvement in multivalent filament binding (Figure 2C). Systematic differential analysis identified 9,954 phosphorylation events with potential functional relevance (Figure 2D) and showed that especially the least soluble third of the proteome is affected: ∼50% of phosphopeptides in this group exhibited significant solubility differences relative to the average protein (moderated t-test, adj. p-value < 0.05, |log₂ fold-change| > log₂(1.5); Figure S2D). On a per-protein basis (for proteins with >3 phosphosites), insoluble proteins were disproportionately affected: nearly 50% of phosphosites showed differential solubility, significantly more than for highly soluble proteins (two-sided unpaired t-test, p-value < 0.05, Figure 2E). As the canonical view of signaling implies that cellular adaptation to perturbations such as BRAF inhibition involves transcriptional and nuclear processes, we complemented the phospho-SPP approach with a subcellular fractionation strategy. Specifically, we performed nuclear enrichment of A2058 cells by differential centrifugation ^27^, followed by proteomic and phosphoproteomic analysis of each fraction (Figure 2F, Supplementary Table 2), to identify phosphorylation events associated with chromatin interaction. The quantified nuclear and non-nuclear (phospho)proteomes corresponded well with known compartment annotations ^28^ (Figure S2E). One example, RELA, a transcription factor known to shuttle between cytoplasm and nucleus, was predominantly non-nuclear at the protein level, while S316-phosphorylated RELA was enriched in the nuclear fraction (Figure 2G). This is consistent with literature linking this site to nuclear translocation ^29^. Similarly, PHLPP1, a TF important in T cell activation, was enriched in the nucleus when phosphorylated at S118, a site recently implicated in nuclear import using an orthogonal methodology ^29^, and now confirmed here (Figure 2H). Systematic comparison of phosphopeptide localization to that of their corresponding proteins using differential analysis revealed 16,161 phosphopeptides potentially impacting nuclear localization (moderated t-test, adj. p-value < 0.05, |log₂ fold-change| > log₂(2), Figure 2I). At a global, per-protein level, we observed that phosphorylation events associated with decreased nuclear localization frequently affect a large proportion of phosphosites on a given protein, whereas increases in nuclear localization upon phosphorylation tend to involve a smaller subset of sites per protein (t-test, p-value < 0.01, Figure 2J). Overall, the two datasets appear to be largely complementary, with 84.5% of phosphosites exhibiting significant changes in only one of the two analyses (Figure S2F). The weak, statistically significant anticorrelation (Figure S2G, t-test on pearson correlation coefficient, p < 0.05) of the signal is likely driven by phosphorylation events with opposing effects on solubility and localization. Examples are nuclear translocation and DNA binding which decreases solubility and increases nuclear localization or mitotic phosphorylation which is associated with nuclear envelope breakdown, leading to increased solubility and decreased nuclear localization (Figure S2H).

**Figure 2:**
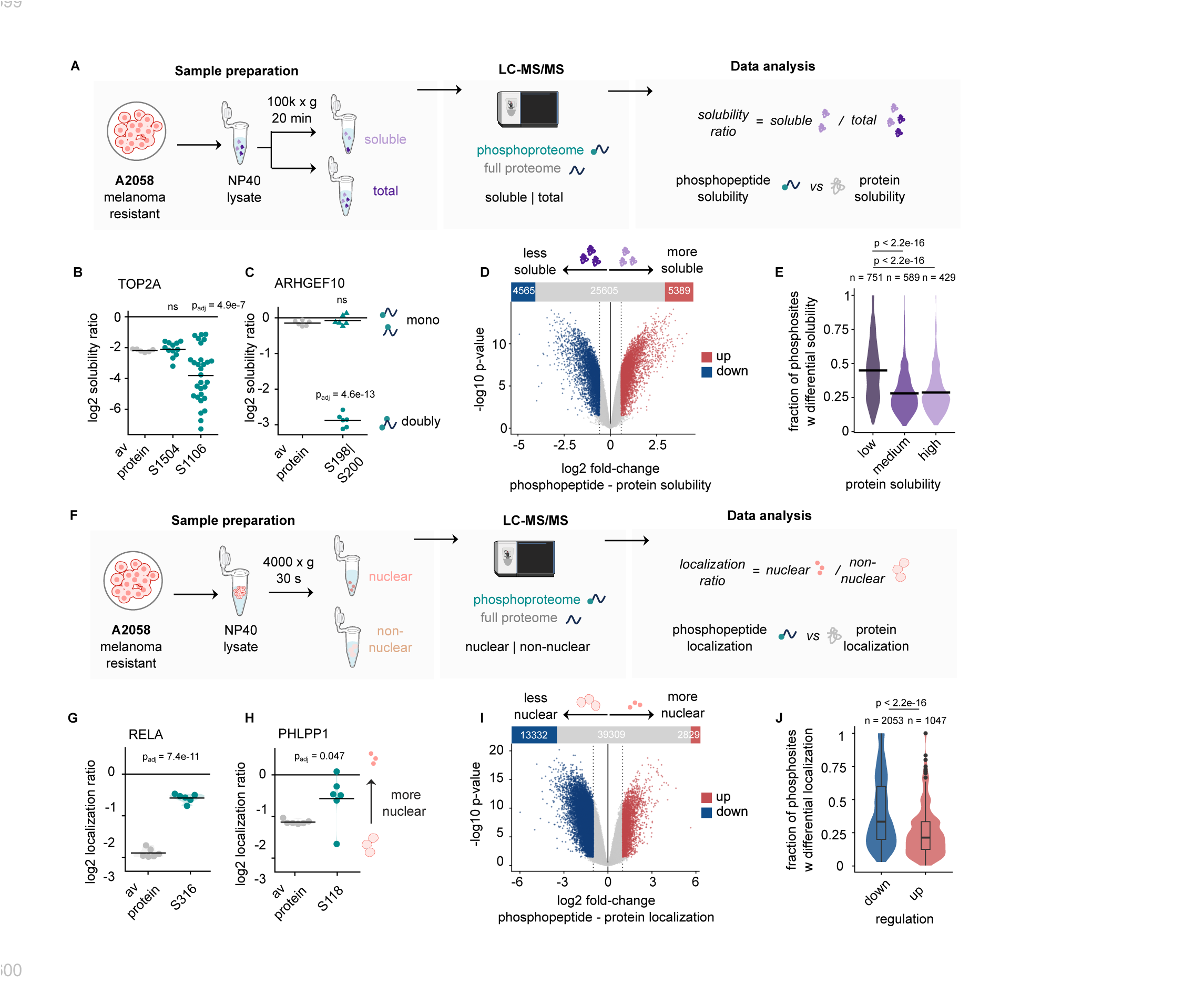
Biophysical phosphoproteomics to investigate aspects of phosphorylation-driven protein regulation. **(A)** Overview of the phosphoproteome-level solubility profiling (phospho-SPP) workflow. A2058 cells were lysed with NP-40 (0.8%), followed by ultracentrifugation (100k g, 20 min) to separate the soluble proteome for comparison to the full proteome. TMT18-based quantitative proteomics was performed with the phosphoproteome and proteome for soluble and total fractions. Solubility was calculated as the ratio of soluble versus total intensities for phosphopeptides and proteins. **(B)** and **(C)** Example solubility profiles for protein (protein intensity in grey) and corresponding phosphoproteoform-specific solubility (cyan): TOP2A (S1504, S1106) and ARHGEF10 (comparison between mono- and doubly phosphorylated states at S198/S200). Significance was assessed using limma’s moderated t-test. **(D)** Volcano plot of differential phosphopeptide solubility relative to protein solubility, indicating phosphopeptides associated with either increased (red) or decreased (blue) solubility compared to the corresponding protein (moderated t-test, adj. p-value < 0.05, |log₂ fold-change| > log₂(1.5) as indicated by dashed line). **(E)** Comparison of the fraction of phosphopeptides showing differential solubility per protein (for proteins with three or more phosphosites) between high, medium and low protein solubility bins (two-sided, unpaired t-test). Black lines indicate the mean. **(F)** Overview of the experimental workflow integrating nuclear fractionation with phosphoproteomics. A2058 cells were fractionated into nuclear enriched and non-nuclear fractions (4000 g, 30 s) followed by TMT18-based phosphoproteomic and proteomic analysis. localization was calculated as the ratio of nuclear versus non-nuclear intensities for phosphopeptides and proteins. **(G)** and **(H)** Example localization profiles for protein (protein intensity in grey) and corresponding phosphoproteoform-specific localization (cyan): RELA (S316) and PHLPP1 (S118). Significance was assessed using limma’s moderated t-test. **(I)** Volcano plot of differential phosphopeptide localization relative to protein localization, indicating phosphopeptides associated with either increased (red) or decreased (blue) nuclear localization compared to the corresponding protein (moderated t-test, adj. p-value < 0.05, |log₂ fold-change| > log₂(2) as indicated by dashed line). **(J)** Comparison of the fraction of phosphopeptides showing differential localization per protein (for proteins with three or more phosphosites) per regulation direction (two-sided, unpaired t-test). Boxplots indicate mean, first and third quartiles.

### Biophysical phosphoproteomics prioritizes functional phosphorylation events in BRAFV600E signaling

We combined the time-resolved phosphoproteomics dataset of Dabrafenib response with the phospho-solubility and localization datasets to investigate the functional impact of deregulated signaling across 64k phosphosites in A2058 cells. This revealed that 31% of Dabrafenib-responsive sites significantly affect protein solubility, nuclear enrichment, or both, thereby markedly reducing the number of potentially relevant features (Figure 3A). The influence of phosphorylation on protein solubility or nuclear enrichment did not correlate with overall treatment response (Figure S3A) showcasing no preferential perturbation of sites that affect biophysical properties. Comparison with computational predictions of phosphorylation functionality ^30^ showed significantly higher predicted functionality for sites exhibiting signal across the biophysical phosphoproteomics datasets (p-value < 0.05, two-sided, unpaired t-test; Figure 3B). While only around 2% of sites have a reported function (Figure 3C), prioritization based on biophysical evidence increases the functional fraction up to 8% (Figure 3D). Of note, the different phosphosite subsets did not differ in regulation strength (log₂ fold-change) in response to Dabrafenib, demonstrating that this prioritization of functionally relevant phosphorylation events cannot be achieved by effect-size filtering (Figure S3B). Secondary-structure predictions of phosphosites using AlphaFold2 ^31^ confirmed that most sites lie within intrinsically disordered regions (IDRs) (Figure 3E). Notably, phosphosites with biophysical evidence were even more enriched in IDRs, occurring less frequently in structured regions than sites without such evidence (Figure 3F). In respect to prior knowledge and annotation depth, the data revealed that around 4% of the phosphosites in the dataset have a literature annotation ^1^ for an upstream kinase (Figure S3C), but this number can increase substantially combining different kinase-substrate prediction resources ^32–34^ (Figure 3G). Phosphosites with biophysical evidence were more likely to have at least one known upstream kinase and, on average, exhibited a higher number of kinase-substrate annotations (Figure 3H, Figure S3D). Taken together the integration of abundance and biophysical phosphoproteomics data prioritizes functionally relevant phosphorylation events of BRAFV600E signaling.

**Figure 3:**
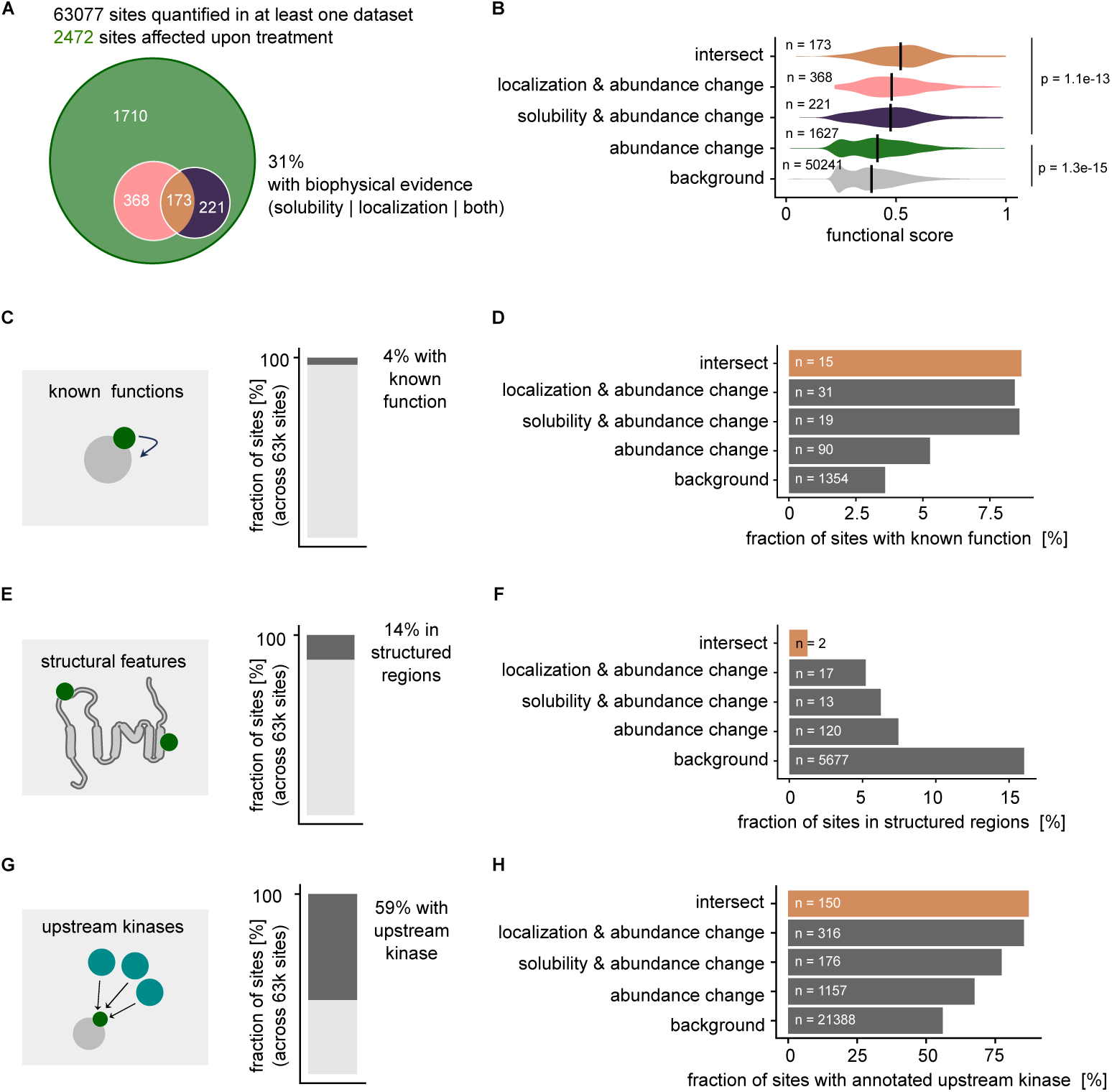
Biophysical phosphoproteomics prioritizes functional phosphorylation events in BRAFV600E signaling. **(A)** Summary of quantified and treatment-responsive phosphosites. Of 63,077 phosphosites quantified in total, 2,472 show Dabrafenib treatment-dependent abundance changes in the A2058 time course experiment. Among these, 31% can be annotated with additional biophysical evidence from solubility changes, localization changes, or both. **(B)** Functional score distributions for different phosphosite groups (as indicated in (A)): abundance-only changes, solubility and abundance changes, localization and abundance changes, and their intersection. Sites with biophysical evidence display significantly higher functional scores than background (two-sided, unpaired t-test, p-value < 0.05). Black lines indicate the mean. **(C)** Fraction of phosphosites with known molecular functions annotated in PhosphoSitePlus ^64^ of all 63k phosphosites quantified across all datasets. **(D)** Fraction of phosphosites with known molecular function within abundance-only, solubility-abundance, localization-abundance, and intersect groups. **(E)** Fraction of phosphosites located in structured regions as predicted by structuremap ^31^ of all 63k sites quantified across all datasets. **(F)** Fraction of phosphosites in structured regions within the same phosphosite groups as in (D). **(G)** Fraction of phosphosites with at least one annotated upstream kinase in an aggregated kinase-substrate network (see methods) of all 63k phosphosites quantified across all datasets. **(H)** Fraction of phosphosites with known upstream kinases in abundance-only, solubility-abundance, localization-abundance, and intersect groups.

### Network-based multi-omics integration to formulate mechanistic hypotheses from multimodal phosphoproteomics data

Given the complexity and connectivity of the BRAFV600E driven phosphorylation events, we next constructed a system-wide model integrating the multimodal phosphoproteomics data with downstream readouts. We leveraged knowledge-driven network modelling to connect and prioritize phosphosites and proteins with high regulatory potential, yielding directly interpretable and experimentally testable hypotheses in the form of multi-omics signaling networks. To capture the molecular downstream consequences of BRAF inhibition, we generated thermal proteome profiling (TPP) ^16^ and time-resolved transcriptomics data in Dabrafenib-treated A2058 cells (Figure S4A, B, C, F, Supplementary Table 3, Supplementary Table 4). TPP provides information on protein thermal stability changes (Figure S4G) as a proxy of altered function ^35^ which can be directly included in models ^36^, whereas transcriptomic analysis enables inference of altered transcription factor activities from changes on their target genes (see methods; Figure S4C). These two modalities reflect complementary consequences of altered phosphorylation on the protein level, encompassing processes ranging from transcriptional regulation to translational control and metabolism (Figure S4E, H, I). We formulated this integrative network analysis as a Rooted Prize-Collecting Steiner Tree (PCST) problem through the framework CORNETO ^37^. In total, we combined ∼2900 nodes, comprising phosphosites, transcription factors, and proteins with altered thermal stability, and ∼13,000 potential edges derived from databases as input for the network optimization (Figure 4A, Figure S5A). Four different input sets were defined by varying phosphosite importance based on the phosphoproteomics measurements (abundance change only, abundance & solubility, abundance & localization, combined weight of all datasets), aiming to construct sets of inputs with different levels of functional evidence. Networks were then built per input set, varying the cost of including nodes to modulate network size, and varying kinase weights to shape the depth of the signaling backbone. In total, 252 networks were generated (Supplementary Table 5), with the largest ones comprising ∼1500 nodes. Across different networks, phosphosites made up ∼80% of network content, underscoring the feasibility of constructing site-level networks with the chosen approach (Figures S5B). Further, the introduction of biophysical evidence selectively affected the inclusion of phosphosites (Figure 4B), as exemplified for the MYLK kinase in networks with and without biophysical evidence (Figure 4C). This highlights the challenge of identifying relevant phosphosites for signaling processes without additional information, as the model’s flexibility allows it to choose among similar sites when relying solely on abundance change estimates. Inclusion of different levels of biophysical evidence did not alter overall network node or edge robustness (Figure S5C,D) or node centralities (Figure S5E), indicating changes in the composition of network nodes rather than in the topology. Independent of the prizing strategy, smaller networks are enriched for phosphosites consistently affected across cell lines (Figure S5F), whereas no trend is observed for the temporal abundance regulation of sites (Figure S5G). Comparing networks built without weighting (abundance only) to those incorporating a combined weight for solubility and nuclear enrichment (combined) revealed that inclusion of biophysical evidence affected around one third of the nodes. To assess the impact of this difference, we performed pathway enrichment analysis across the networks (p-value < 0.05, normalized weighted mean test). Pathways related to BRAFV600E signaling, including MAPK signaling, MYC regulation, transcription, proliferation, metabolism, and phosphorylation, showed significantly stronger enrichment in the networks with additional weighting (two-sided Fisher’s exact test, p-value < 0.05, Figure 4D), suggesting that integration of biophysical evidence increases the biological relevance of the resulting models.

**Figure 4:**
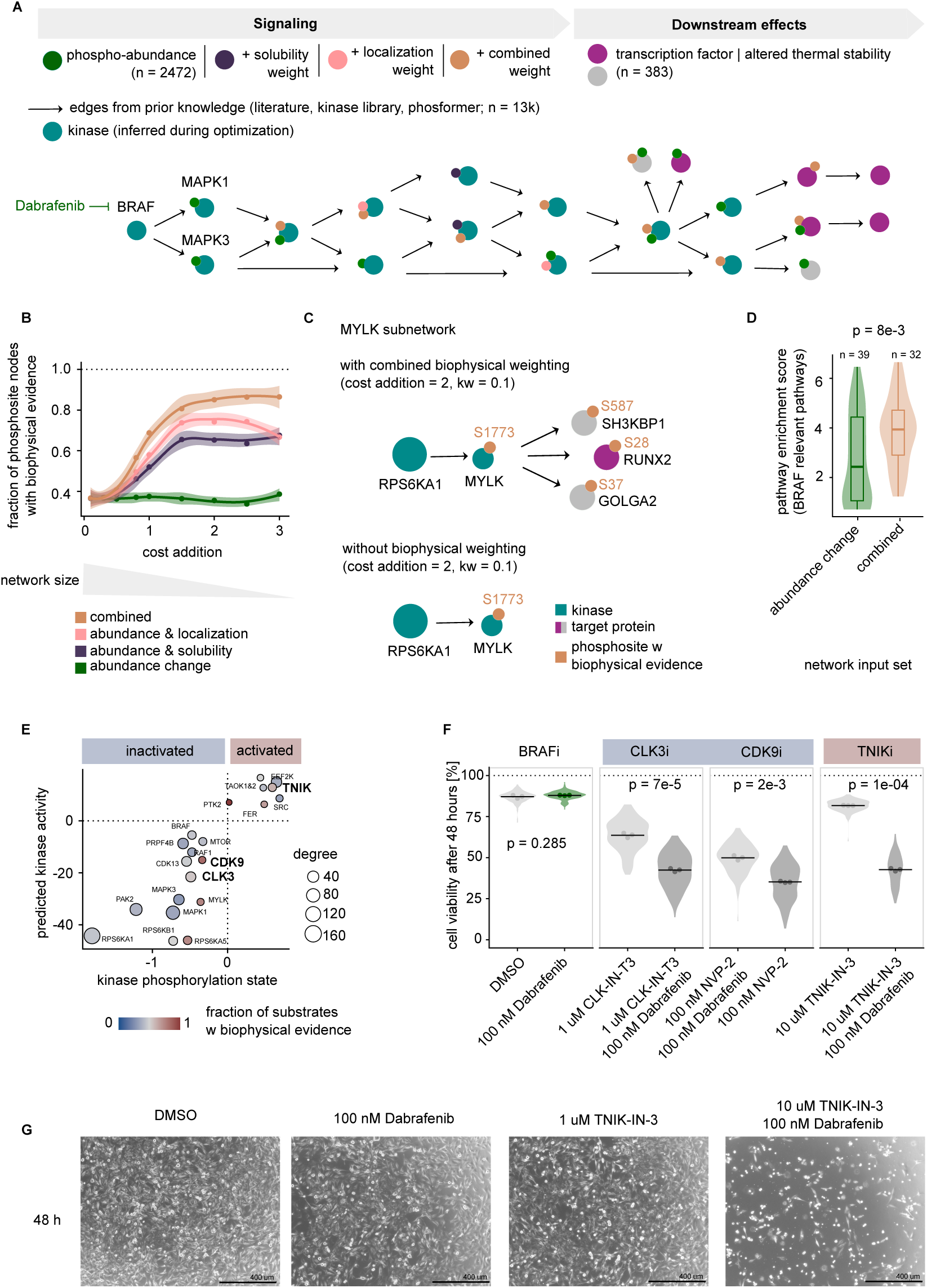
Network-based multi-omics integration to formulate mechanistic hypotheses from multimodal phosphoproteomics data. **(A)** Conceptual overview of the multi-omics network modeling approach. Multimodal phosphoproteomics, transcriptomics and thermal proteome profiling (TPP) data was combined to optimized subnetworks of directed signaling downstream of BRAFV600E. Networks were modeled with different weights (abundance-only changes, solubility-abundance changes, localization-abundance and their combination) to study the impact of biophysical evidence on signaling networks and different cost additions and kinase weights to vary size and depth. **(B)** Fraction of phosphosite nodes with biophysical evidence (solubility, localization, or both) recovered across network sizes and applied weighting strategy, shown as a function of cost addition.**(C)** Example subnetwork for the MYLK kinase constructed with combined biophysical weighting or only based on abundance changes for a cost addition of 2 and a kinase weight of 0.1. Top: Combined biophysical weighting expands the local subnetwork and identifies additional MYLK target sites with solubility or localization changes (S587 on SH3KBP1, S28 on RUNX2, S37 on GOLGA2). Bottom: Without biophysical weighting, the subnetwork is smaller and includes fewer regulated substrate candidates. Colours indicate kinase, target protein, and phosphosites with biophysical evidence. **(D)** Pathway enrichment scores for BRAF-relevant pathways comparing networks reconstructed using abundance-change information alone versus using combined biophysical weighting (normalized weighted mean test, p-value < 0.05, selected top pathways). Boxplots indicate mean, first and third quartiles. **(E)** Predicted kinase activities for kinases in the network model plotted against kinase phosphorylation states measured in the Dabrafnib time course (filtered for kinases with correlated signal). Colour indicates the fraction of substrates with biophysical evidence across all network models with combined biophysical weighting. Point size indicates mean degree centrality (= number of substrates) across networks. **(F)** Cell viability assays in A2058 cells quantifying SYTOX fluorescence following inhibition of selected kinases predicted to be functionally relevant (CLK3, CDK9, TNIK) for 48 hours (unpaired, two-sided t-test). The experiment was performed in three biological replicates (points), 56 data points were generated per replicate and condition (violin shape). Black lines indicate the mean. **(G)** Representative images of A2058 cells treated for 48 hours with DMSO, Dabrafenib, TNIK-IN-3 or their combination. Scale bars: 400 μm.

### Network models identify novel kinase targets and phosphorylation-linked transcriptional control

Having integrated the different site and protein level data in a network model, we next leveraged it to identify kinases relevant for BRAFV600E-driven signaling strongly supported by biophysical evidence, with the goal of validating their functional relevance experimentally. We first calculated the fraction of substrates with biophysical evidence across all network solutions built with combined biophysical evidence and compared this to the overall degree (= number targets across networks) of the kinase in the networks. (Figure S5H). Alongside expected kinases implicated in melanoma signaling such as PRKD3 ^38^, RPS6KA5 ^39^, and ROCK2 ^40^, this analysis highlighted poorly characterised candidates including TNIK, CDK9 and CLK3. The subsequent kinase activity inference (p-value < 0.05, normalized weighted mean test) and phosphorylation analysis supported these observations as predicted CDK9 and CLK3 activities were reduced upon Dabrafenib treatment, while predicted TNIK activity was increased and accompanied by upregulated phosphorylation (Figure 4E, Figure S5I). In order to validate the relevance of these kinases for melanoma signaling, we tested combination treatments of Dabrafenib with selective inhibitors (NVP2, CLK-IN-T3, TNIK-IN-3) and assessed their impact on cell viability. Co-targeting BRAF and these kinases significantly reduced cell survival (p-value < 0.05, two-sided, unpaired t-test; Figure 4F, Figure 4G). In line with the predicted activities of these kinases, for CDK9 and CLK3 inhibition, a more synergistic effect was observed, whereas Dabrafenib and TNIK co-inhibition produced a synthetic lethal-like effect. Importantly, Dabrafenib and TNIK co-inhibition enhanced cytotoxicity not only in the A2058 cells but also in the two additional BRAFV600E-mutant models (MNT-1, COLO-201), indicating general relevance of this signaling axis (Figure S5J). Collectively, these findings highlight the power of the network model to reveal kinases functionally involved in BRAFV600E signaling, including both established components and newly implicated regulators such as TNIK, CDK9, and CLK3.

### ETV3 activity is phospho-regulated in response to BRAF inhibition across cell lines

While kinase signaling is a central aspect of the network models, they also provide a rich contextual resource to investigate individual phosphorylation events, including predicted upstream kinases, site-specific functional effects, and potential downstream consequences. The models propose mechanistic hypotheses for up to 1200 phosphosites, many of which previously lacked functional annotations or known upstream kinases in literature (Figure S6A). Interesting examples include phosphorylation events on transcriptional regulators which are central for the long-term response to the BRAF inhibition such as MYC, FOS, and JUN, positioned downstream of MAPK1 in the network models (Figure 5A). Detailed inspection of MYC in the network models revealed five phosphosites downregulated upon Dabrafenib treatment (S62|T58, S281|S293, S344 prioritized out of ten sites deregulated in total; Figure 1H), while the transcriptomics data based activity prediction suggested MYC inactivation after 4 hours (Figure S6B). Regarding the biophysical phosphoproteomics data, S62 and T58, which are known to functionally interact ^21^, as well as S281 and S293 whose interaction is reported but less characterized ^41^, showed differential solubility and localization for the mono and double phosphorylated forms (Figure S6C and S6D). Besides these major and known oncogenic transcription factors, the network analysis also uncovered previously unknown regulators of BRAFV600E signaling such as the transcriptional repressor ETS translocation variant 3 (ETV3, also called PE-1 or METS; Figure 5B). While phosphorylation of ETV3 by MAPK1 has been reported ^42^, its involvement in BRAF-driven signaling has not been described previously. Dabrafenib treatment led to dephosphorylation of ETV3 at three sites within 30 minutes of treatment (Figure 5B), including S29 and S139, which were prioritized by the network model. Transcription factor activity analysis predicted strong ETV3 activation within two hours of Dabrafenib treatment (Figure 5C). In the biophysical phosphoproteomics data, ETV3 appeared highly insoluble (Figure 5D), consistent with its reported nuclear localization and DNA-bound state ^42^. The S29-phosphorylated form was significantly more soluble than the average protein (moderated t-test, adjusted p-value < 0.05, |log₂ fold-change| > log₂(1.5), Figure 5D), whereas the S139 phosphopeptide could not be detected in this dataset. In the TPP data, the unmodified counterpart of the S139 site exhibited increased thermal stability compared to other ETV3 peptides (Figure 5E). Both prioritized sites are located in close proximity to the Ets DNA-binding domain of ETV3 (Figure 5F). Together with the biophysical data this suggests a functional role for these sites in regulating ETV3 DNA binding. Notably, Dabrafenib treatment increased the insoluble fraction of ETV3, consistent with enhanced chromatin association upon dephosphorylation (Figure 5G). To further collaborate the link between solubility and DNA-binding, pellets from Dabrafenib-treated cells were subjected to DNase digestion and ETV3 solubility was assessed subsequently. DNA digestion resolubilized ETV3, indicating that the Dabrafenib-induced solubility decrease of ETV3 is DNA-dependent (Figure 5H). Of note, ETV3 dephosphorylation upon Dabrafenib treatment occurs across BRAFV600E-mutant cell lines (A2058, MNT-1, COLO-201; Figure S6E), indicating that ETV3 regulation by mutant BRAF could be context-independent. Further, steady-state phosphoproteomics data generated for a panel of cell lines (Figure S6F, G) showed that hyperphosphorylated ETV3 was more abundant in BRAFV600E-mutant compared to non-mutant cells (p-value < 0.05, two-sided, unpaired t-test; Figure S6H, Supplementary Table 2). Taken together, these findings suggest that ETV3 activation upon BRAF inhibition is mediated by rapid dephosphorylation, which enhances DNA binding and thus that ETV3 hyperphosphorylation is repressing its activity in BRAFV600E-mutant cells.

**Figure 5:**
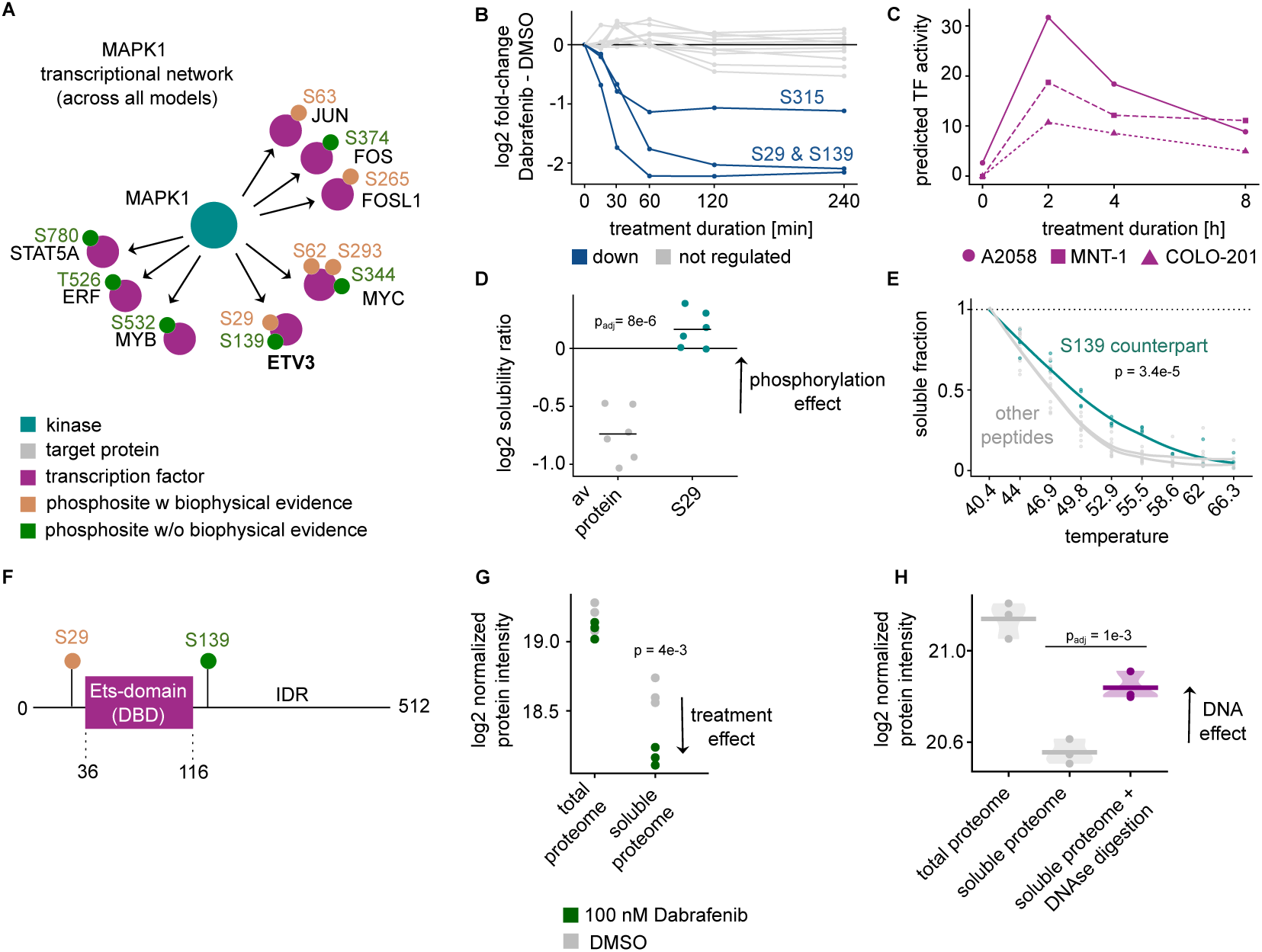
Network models identify novel kinase targets and phosphorylation-linked transcriptional control. **(A)** MAPK1-centred transcriptional subnetwork aggregated across all generated network models. Nodes represent transcription factors, kinases, and target proteins; edges show inferred regulatory relationships. Phosphosites with solubility or localization evidence are highlighted. ETV3 exhibits two MAPK1-regulated, potentially functional sites (S29 and S139). **(B)** Temporal phosphorylation dynamics of ETV3 following Dabrafenib treatment (100 nM, 4 h) in A2058 cells. Significance was assessed using limma’s moderated t-test. **(C)** Predicted transcription factor (TF) activity of ETV3 across three BRAFV600E-mutant cell lines (A2058, MNT-1, COLO-201). **(D)** log₂ solubility ratios for the average ETV3 protein and the S29 phosphoproteoform across six replicates. Significance was assessed using limma’s moderated t-test. **(E)** Thermal stability profiles for the S139 peptide (unmodified counterpart of the phosphopeptide) versus other ETV3 peptides. Significance was assessed using an unpaired, two-sided t-test. **(F)** Schematic of the ETV3 structure. Indicated are two prioritized phosphosites (S29 and S139) as well as the DNA binding domain (DBD) of ETV3 which is the only structured part of the protein. **(G)** log₂ ETV3 protein intensity in total or NP40-soluble lysis conditions for three replicates for DMSO and Dabrafenib (100 nM, 4 hours) treated cells. Significance was assessed with an unpaired, two-sided t-test. **(H)** log₂ ETV3 protein intensity in total, NP40-soluble and NP40-soluble combined with DNAse digestion lysis conditions for three replicates for Dabrafenib (100 nM, 4 hours) treated cells. Significance was assessed using limma’s moderated t-test.

### ETV3 activation leads to a druggable metabolic vulnerability in BRAFV600E-mutant cells

To evaluate the functional role of ETV3 downstream of BRAF inhibition, we performed time-resolved quantitative proteomics in ETV3 knockdown cells treated with Dabrafenib (Figure 6A, Figure S7A, B, Supplementary Table 6). The proteome-wide remodeling upon ETV3 knockdown and Dabrafenib treatment is consistent with the model prediction of ETV3 activation by BRAF inhibition (Figure 6B). ETV3 knockdown altered the expression of 684 proteins upon BRAF inhibition, clustering into early direct changes and later secondary effects (Figure 6C, Figure SC). Among proteins expressed in an ETV3-dependent manner, Glucose transporter type 3 (GLUT3 or SLC2A3) stood out as its expression during Dabrafenib was strongly dependent on ETV3 knockdown (Figure 6D) and transcriptionally repressed in cells with ETV3 present (Figure S7D). This data links ETV3 activity to glucose metabolism, since GLUT3 is a high-affinity, insulin-independent glucose transporter relevant for cellular growth in low glucose conditions ^43^. It is also noteworthy that GWAS results ^44,45^ associate single-nucleotide polymorphisms (SNPs) at the ETV3 gene locus with a spectrum of glucose- and immune-related phenotypes and diseases at the population level (Figure S7G). ETV3 knockdown combined with Dabrafenib treatment and metabolomics of culture supernatants confirmed the role of ETV3 in glucose metabolism. Upon ETV3 knockdown, cells took up significantly more glucose than control cells when treated with Dabrafenib (p-value < 0.05, two-sided, unpaired t-test; Figure 6E, Figure S7E). Both conditions (Dabrafenib and DMSO) showed increased glutamine uptake, but this was significantly stronger in control cells (p < 0.05, two-sided, unpaired t-test), suggesting that ETV3 promotes a shift towards glutamine dependency in BRAFV600E cells upon BRAF inhibition (Figure 6E). The observed metabolic consequences of BRAFV600E-driven signaling and BRAF inhibitor resistance suggest that A2058 cells would be susceptible to disruptions in energy metabolism in combination with BRAF inhibition. To test this, we combined Dabrafenib treatment with the glutaminolysis inhibitor Telaglenastat, which revealed a potential synthetic lethal interaction between BRAF and GLS activity in A2058 cells. (Figure 6F, Figure 6G). Importantly, this combinatorial vulnerability was also observed in COLO-201 and MNT-1 cells with varying magnitude, underlining its broader relevance also in cells sensitive to BRAF inhibition (Figure S7F). Sensitivity to the combination correlated with the extent of GLUT3 downregulation upon Dabrafenib treatment (Figure S7D). Together, these results establish that ETV3 is a phosphoregulated transcriptional repressor activated by BRAF inhibition, with broad effects including rewiring cellular metabolism. Mechanistically, ETV3 represses GLUT3 expression and drives melanoma cells towards glutamine dependency given BRAF inhibition, creating a vulnerability that can be exploited therapeutically. This example highlights how systematic biophysical phosphoproteomics not only enables construction of mechanistic models but also allows extraction of actionable therapeutic hypotheses, providing both mechanistic insight and potential biomarkers for treatment response.

**Figure 6:**
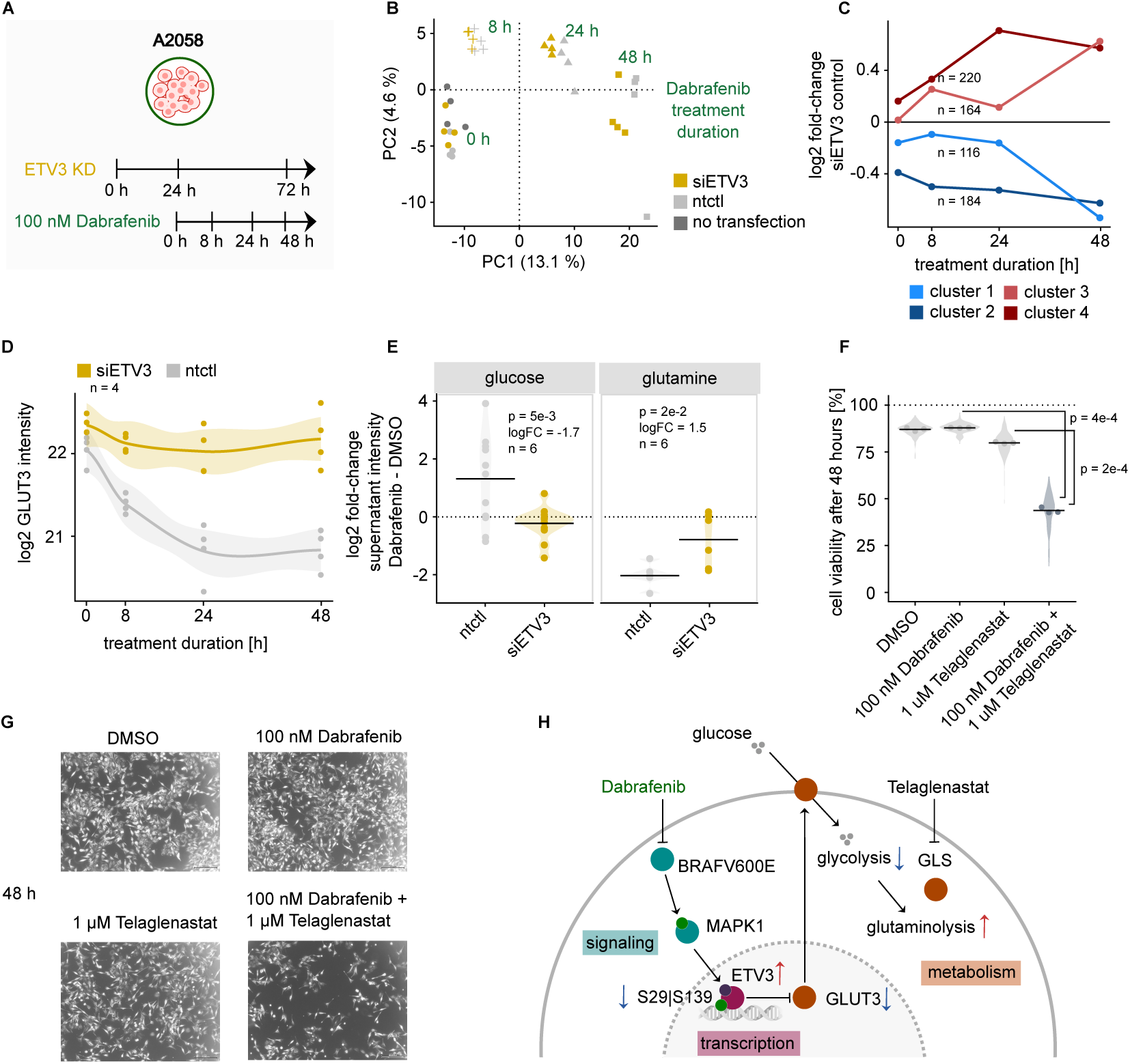
ETV3 activation leads to a druggable metabolic vulnerability in BRAFV600E-mutant cells. **(A)** Design overview for ETV3 follow-up experiments. A2058 melanoma cells underwent ETV3 knockdown (siETV3) or non-targeting control (ntctl), followed by Dabrafenib treatment (100 nM) monitored by time-resolved quantitative proteomics and metabolomics. **(B)** PCA of the protein intensities in A205 cells during Dabrafenib treatment (shape, times indicated) and ETV3 knockdown (colour). The PCA was calculated based on the top 10% most variable proteins across samples. **(C)** Time-resolved log₂ fold changes comparing protein intensities in siETV3 knockdown cells to control cells, grouped into four expression clusters based on their variation during Dabrafenib treatment. **(D)** GLUT3 (SLC2A3) protein abundance across the Dabrafenib time course in siETV3 knockdown cells and control cells. Fit lines indicate mean protein intensities (loess fit). **(E)** Metabolomics quantification of glucose and glutamine intensity in cellular supernatant following Dabrafenib treatment and ETV3 knockdown. ETV3 knockdown increases glucose uptake (left) and decreases glutamine uptake (right) compared to the control given Dabrafenib treatment. Black lines indicate the mean. **(F)** Cell viability after 48 hours under single or combined treatments. Alone, Dabrafenib (100 nM) or the glutaminase inhibitor Telaglenastat (1 μM) have limited effects, whereas the combination markedly reduces viability (unpaired, two-sided t-test). The experiment was performed in three biological replicates (points), 56 data points were generated per replicate and condition (violin shape). Black lines indicate the mean. **(G)** Representative microscopy images of A2058 cells after 48 hours of treatment. Scale bars: 400 μm. **(H)** Model summarizing the proposed mechanism of ETV3 regulation. BRAFV600E-driven MAPK signaling phosphorylates and regulates ETV3 (S29, S139), which in turn controls GLUT3 expression Dabrafenib reduces MAPK signaling and ETV3 phosphorylation, decreasing glucose uptake leading to metabolic collapse and cell death when combined with the glutaminolysis inhibitor Telaglenastat.

## Discussion

Hyperactive kinases such as mutant BRAF induce system-wide effects requiring complex signaling adaptations to sustain cellular viability in treatment resistant models^12^. We present a comprehensive model of BRAFV600E-driven signaling combining phosphoproteomics, biophysical proteomics, transcriptomics, and computational modeling which allowed us to identify novel strategies to resensitize resistant melanoma cells to BRAF inhibition. Our phosphoproteomic analyses revealed that while a shared set of phosphorylation events defines the core BRAF response across BRAFV600E-mutant cell lines, most signaling changes are cell line-specific, illustrating that cells with the same oncogenic driver mutation adopt different signaling states (Figure 1). Time-resolved data reveals the broad immediate impact of BRAF inhibition highlighting the difficulty in assigning linear pathway logic to complex cellular systems ^46^. While these phosphoproteomic data outline how BRAF inhibition affects cellular signaling, the impact on protein function of the vast majority (96%) of these changes are unknown ^14^. We applied biophysical proteomics on the phosphopeptide level measuring solubility and localization changes as proxies for phosphorylation-dependent protein regulation (Figure 2). This provides an experimental and unbiased complement to computational prediction ^30^ or genetic perturbation-based ^29,47^ strategies ^17,19^. Integrating these datasets with the BRAF inhibition data revealed measurable biophysical signals for around 1/3 of Dabrafenib-affected phosphosites, (Figure 3). The level of measured biophysical evidence is correlated with predicted functional scores for phosphorylation sites ^30^. Notably, biophysical evidence was independent of regulation strength indicating that these approaches capture orthogonal, unbiased information on phosphosite relevance. To place these site-level findings into a broader mechanistic context, we developed an optimisation-based network modelling framework that links phosphorylation events to downstream functional outcomes (Figure 4). Integration of biophysical evidence affects network composition and significantly improves the signal for BRAF related pathways in the network and enabled us to identify kinases outside the canonical MAPK pathway, including TNIK, CDK9, and CLK3, whose co-inhibition with BRAF efficiently resensitised A2058 cells to treatment. The network further provides phosphosite-level mechanisms, capturing both well-established regulators (e.g., MYC S62-T58, FOS S374, JUN S63) and previously uncharacterized candidates linked to BRAFV600E signaling. We focused on ETV3, a transcriptional repressor prioritized through the network model. Previous studies linked ETV3 to MAPK signaling and macrophage differentiation ^42,48–53^, but, to our knowledge, no connection to BRAFV600E-driven processes has been reported. Our data suggested that BRAF inhibition promotes ETV3 activation through dephosphorylation on S29 and S139 followed by enhanced chromatin association indicating that phosphorylated forms are less DNA-bound (Figure 5), consistent with data for ETS-domain-containing transcription factors ^42,54^. Functionally, ETV3 influences metabolic rewiring upon BRAF inhibition, reflected by ETV3-dependent repression of the glucose transporter GLUT3 which results in decreased glucose and increased glutamine uptake (Figure 6). Given that GLUT3 is a high-affinity glucose transporter typically expressed in neurons ^43^, its induction in BRAFV600E-mutant cells likely provides a growth advantage potentially driven by ETV3 inactivation through hyperphosphorylation. Accordingly, combined inhibition of BRAF and glutaminolysis sensitized A2058 cells to treatment, revealing a potential metabolic vulnerability. The consistent regulation of ETV3 across BRAFV600E-mutant lines suggests that this mechanism could be broadly relevant, positioning ETV3 as a mediator linking oncogenic signaling to metabolic rewiring which is increasingly recognised as central to cancer cell survival ^55,56^. This is further supported by GWAS data ^44,45^ that associate genetic variation at the ETV3 locus beyond the reported immune regulatory function ^48,52^ with glucose metabolism-related traits and diabetes, a disease characterized by dysregulated glucose transporters ^57^. Although phosphoproteomic and transcriptomic analyses have individually provided valuable insights into BRAFV600E-mutant signaling ^58,59^, an integrated, multimodal strategy that combines in-depth quantitative and biophysical phosphoproteomics with transcriptomics and thermal proteome profiling has not yet been used to construct causal models of this process. Here, this approach is used to suggest mechanisms of drug synergy which is particularly relevant in the context of current clinical practice. BRAF inhibitors represent first-line therapies for BRAF-mutant cancers but are administered in combination to mitigate paradoxical MAPK activation ^60^ and improve response durability. At present, the only approved combinations pair BRAF inhibitors with MEK inhibitors, which are associated with considerable toxicity and target the same signaling pathway in close proximity increasing the risk of resistance emergence ^61^. We show inference of potential drug combinations for a selection of kinases and for ETV3, but in principle, the multi-omics network and biophysical phosphoproteomics highlight more unexplored regulators of BRAFV600E signaling. In general, this approach can be extended to other oncogenic kinases or targeted therapies, enabling systematic investigation of cancer signaling networks at scale. Moreover, applying this strategy across diverse cell or tissue contexts could delineate to which extent functional signaling is conserved or context-dependent. Overall, the framework offers a generalizable strategy to move beyond descriptive phosphoproteomics toward a mechanistic, systems-level understanding of signaling functionality.

## Limitations

- **Stochastic sampling:** The inherent stochasticity of peptide detection in bottom-up MS constrains site overlap across datasets and replicates. In this study, we mitigate this limitation by generating multiple ultradeep datasets, a strategy that, with continued technological advances, will hopefully become increasingly accessible.
- **Phosphoproteoforms**: Distinguishing between differentially modified phosphoproteoforms remains difficult in bottom-up approaches, complicating interpretation of site-specific regulation.
- **Indirect measurement:** Changes in solubility or subcellular distribution provide indirect proxies for potential protein functions and therefore must be interpreted with caution. This approach as well as the network modeling should be viewed primarily as a powerful approach for hypothesis generation rather than as definitive causal evidence.
- **Broader functional potential:** Additional, unexplored functions of ETV3 beyond the metabolic angle examined here are likely and may also contribute to the observed phenotypic effects. The datasets generated in this study provide an ideal starting point for uncovering these further roles.

## Author contributions

M.L.B.: Conceptualization, Data Curation, Formal Analysis, Investigation, Methodology, Project Administration, Software, Visualization, Writing – Original Draft Preparation; M.G.R.: Conceptualization, Software, Supervision, Writing – Original Draft Preparation, Writing – Review & Editing; P.A.R.M.: Conceptualization, Investigation, Supervision, Writing – Review & Editing; D.P.: Investigation, Methodology; I.B.: Investigation, Methodology; D.M.S.: Investigation, Methodology; C.P.: Investigation, Methodology; F.J.: Investigation, Methodology; M.Z.: Conceptualization, Supervision, Funding Acquisition, Resources, Writing – Review & Editing; J.S.R.: Conceptualization, Supervision, Funding Acquisition, Resources, Writing – Review & Editing; M.M.S.: Conceptualization, Supervision, Funding Acquisition, Resources, Writing – Review & Editing

## Acknowledgements

We gratefully acknowledge the invaluable support of several EMBL core facilities and colleagues. We thank the EMBL Proteomics Core Facility, particularly Mandy Rettel and Jennifer Schwarz for operational and experimental assistance, and Frank Stein for help with data handling and processing. We also thank the EMBL Genomics Core Facility, especially Vladimir Benes, for their expertise in transcriptomics. For advice on data analysis and modelling, we thank Stephan Gade and Pablo Mier-Rodriguez. We further thank Nadine Tüchler and Christine Rummel for assistance with transcriptomics data generation. We thank Sophie Moggridge and Lena Müller for great help with the manuscript and Elias De Lamo Peitz and Yuki Hayashi for the support with experiments. Finally, we are grateful to all members of the Savitski Team and the PCF for their extensive and insightful discussions and feedback.

## Conflicts Of Interest

JSR reports funding from GSK, Pfizer and Sanofi and fees/honoraria from Travere Therapeutics, Stadapharm, Astex, Pfizer, Vera Therapeutics, Grunenthal, Tempus, Moderna and Owkin.

## Figures

**Figure S1.**
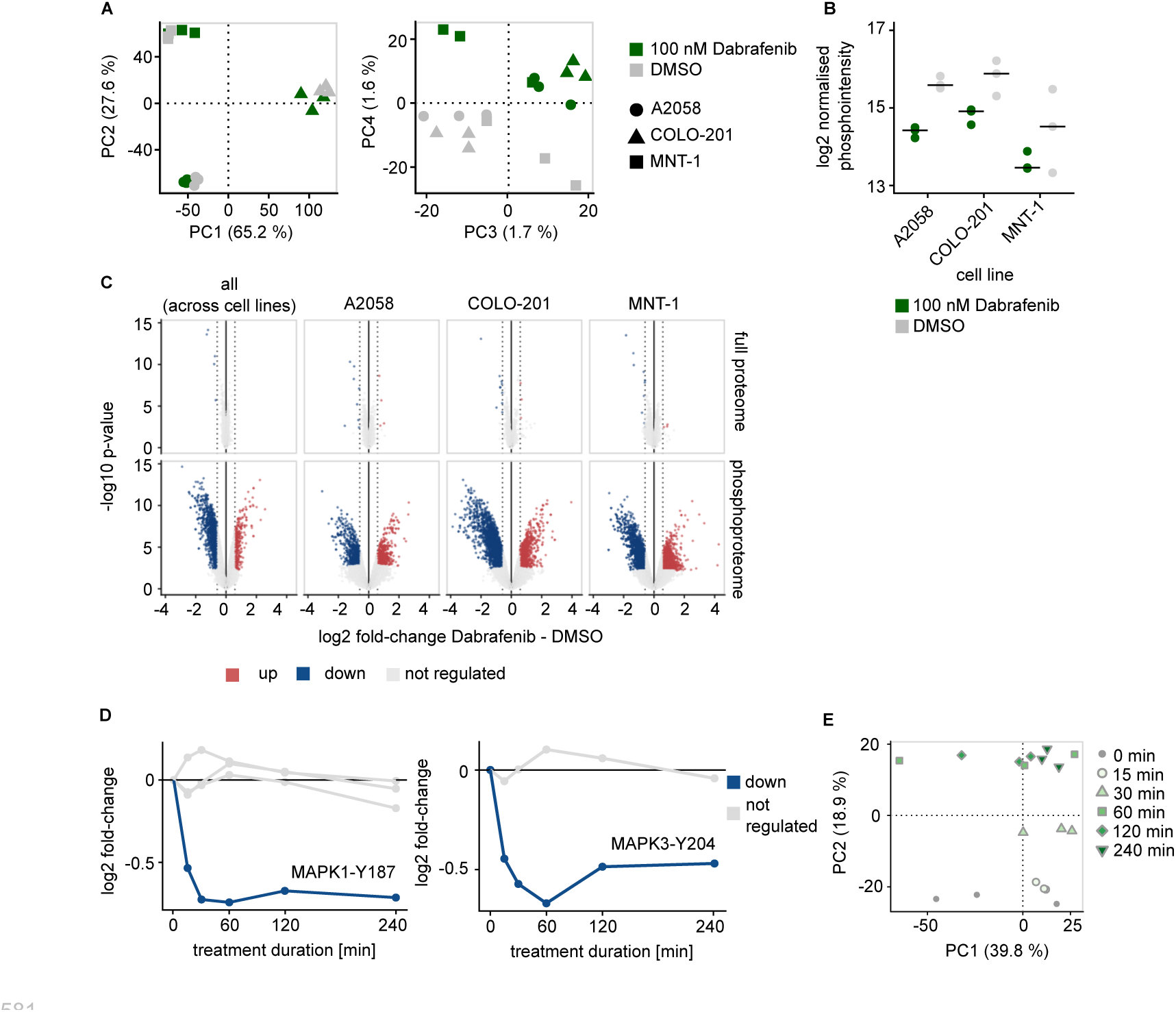
**(A)** Principal component analysis (PCA) of normalized phosphopeptide intensities in A2058 (circle), COLO-201 (triangle), and MNT-1 (square) cells treated with 100 nM Dabrafenib (green) or DMSO (grey) for 4 hours. The PCA was calculated using the top 10% most variable phosphopeptides across samples. **(B)** Normalized MYC S62 phosphorylation (KFELLPTPPLpSPSR) across the three cell lines in response to 100 nM Dabrafenib (4h, green) or DMSO (4h, grey). **(C)** Volcano plots showing differential protein (upper row) or phosphopeptide (lower row) abundance upon Dabrafenib treatment (100 nM, 4 h) relative to DMSO for each cell line or across cell lines (all, interaction effect model). Significantly upregulated (red) and downregulated (blue) features are highlighted (moderated t-test, adj. p-value < 0.05, |log₂ fold-change| > log₂(1.5) as indicated by dashed line). **(D)** Time-resolved phosphorylation dynamics of MAPK1 (Y187, VADPDHDHTGFLTEpYVATR) and MAPK3 (Y204, IADPEHDHTGFLTEpYVATR) activation loop sites, upon Dabrafenib treatment. Significantly downregulated (blue) phosphopeptides are highlighted (moderated t-test, adj. p-value < 0.05, |log₂ fold-change| > log₂(1.5)). **(E)** PCA of normalized phosphopeptide intensities in the Dabrafenib time course (100 nM) across six timepoints (indicated by colour and shape). The PCA was calculated using the top 10% most variable phosphopeptides across samples.

**Figure S2.**
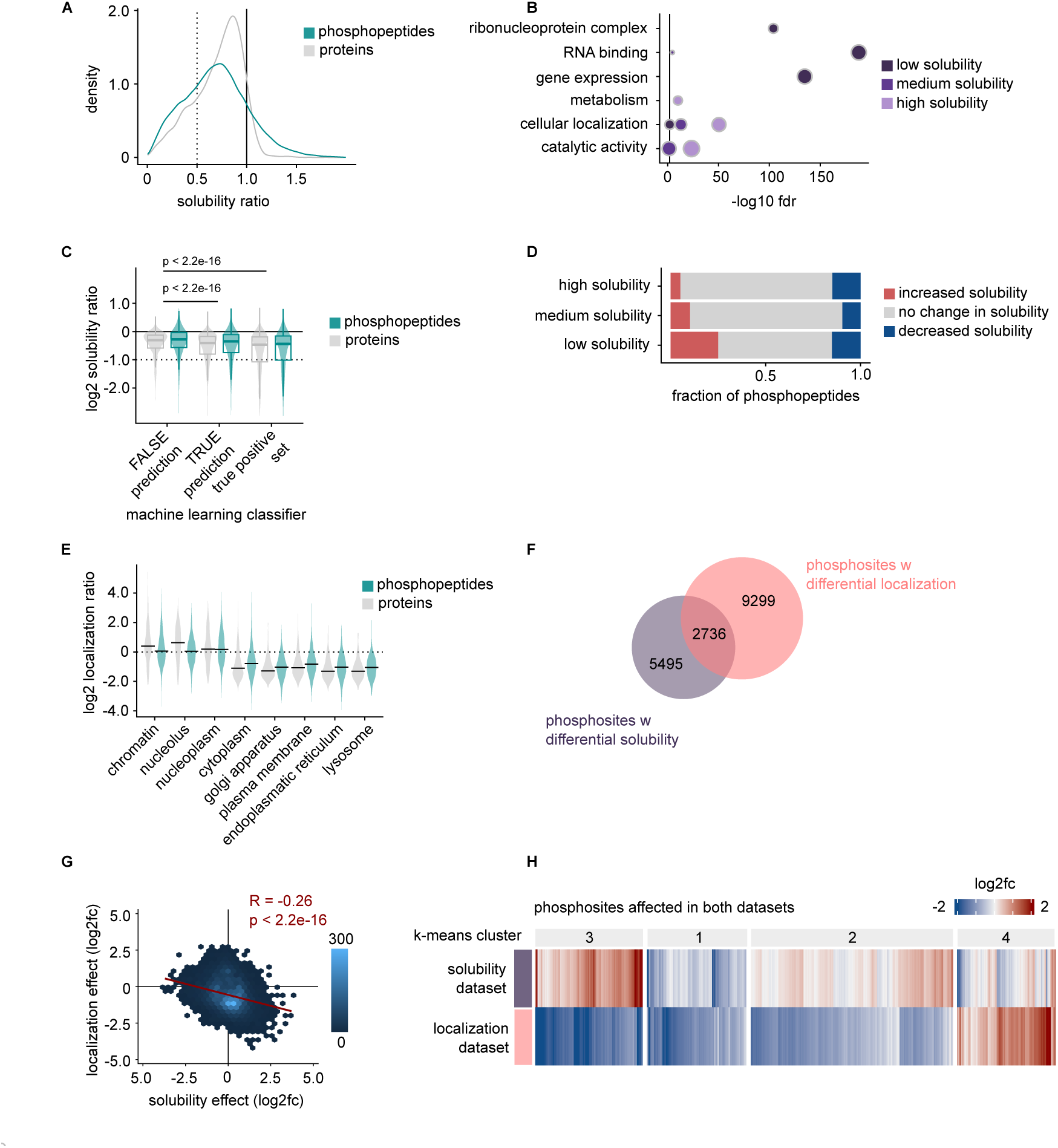
**(A)** Distribution of solubility ratios (= soluble intensity / total intensity) for the full proteome (grey) and phosphoproteome (cyan). **(B)** Gene ontology enrichment analysis of proteins grouped by solubility bins (low, medium, high). Dot size reflects the number of genes per category; colour indicates solubility bin (overrepresentation test, fdr < 0.05). Black line indicates fdr cutoff. **(C)** Comparison of solubility ratios between the full proteome (grey) and phosphoproteome (cyan) per model classifier groups of Hadarovich et al ^63^ which predict condensation probability for proteins. Boxplots indicate mean, first and third quartiles. **(D)** Fraction of phosphopeptides with increased (red), decreased (blue), or not altered (grey) solubility compared to the average protein, stratified by protein solubility class. **(E)** log₂ localization ratios (= nuclear intensity / non-nuclear intensity) of full proteome (grey) and phosphopeptide (cyan) measurements for different compartment annotations of the COMPARTMENTS database ^28^. black lines indicate the mean. **(F)** Overlap between phosphosites showing differential solubility and localization relative to corresponding protein measurements. **(G)** Pearson correlation (R = −0.26, p-value < 2.2e−16) of phosphopeptide solubility and localization effects (phosphopeptide-protein differences) averaged to mean log₂ fold-change per site for sites which are significantly affected in at least one dataset.**(H)** Heatmap of phosphosite effects (mean log₂ fold-change per site) for sites affected in the solubility and localization dataset.

**Figure S3.**
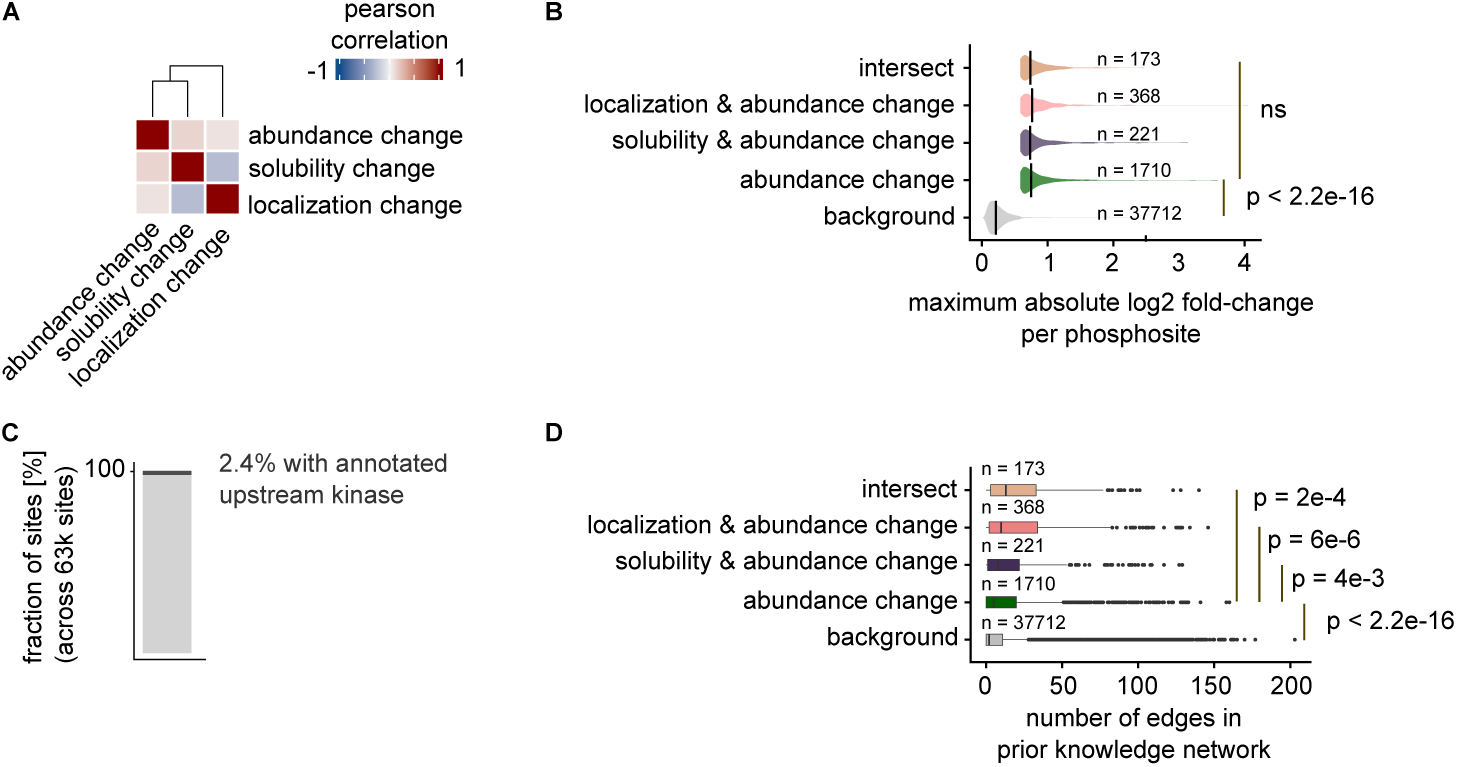
**(A)** Heatmap of pearson correlation of log₂ fold changes across different phosphosite groups and datasets: abundance-altered (Dabrafenib-DMSO), solubility-affected and localization-affected (protein-phosphopeptide). **(B)** Maximum absolute log₂ fold-change of phosphosites in the Dabrafenib time course data compared for sites in abundance-only changes, solubility-abundance changes, localization-abundance and their intersect groups (two-sided, unpaired t-test). Black lines indicate the mean. **(C)** Fraction of phosphosites with supporting evidence from literature resources (PhosphositePlus) of all 63k phosphosites quantified across all datasets. **(D)** Comparison of the number of edges in an aggregated kinase-substrate network (see methods) for phosphosites in abundance-only changes, solubility-abundance changes, localization-abundance and their intersect groups (unpaired, two-sided t-test). Boxplots indicate mean, first and third quartiles.

**Figure S4.**
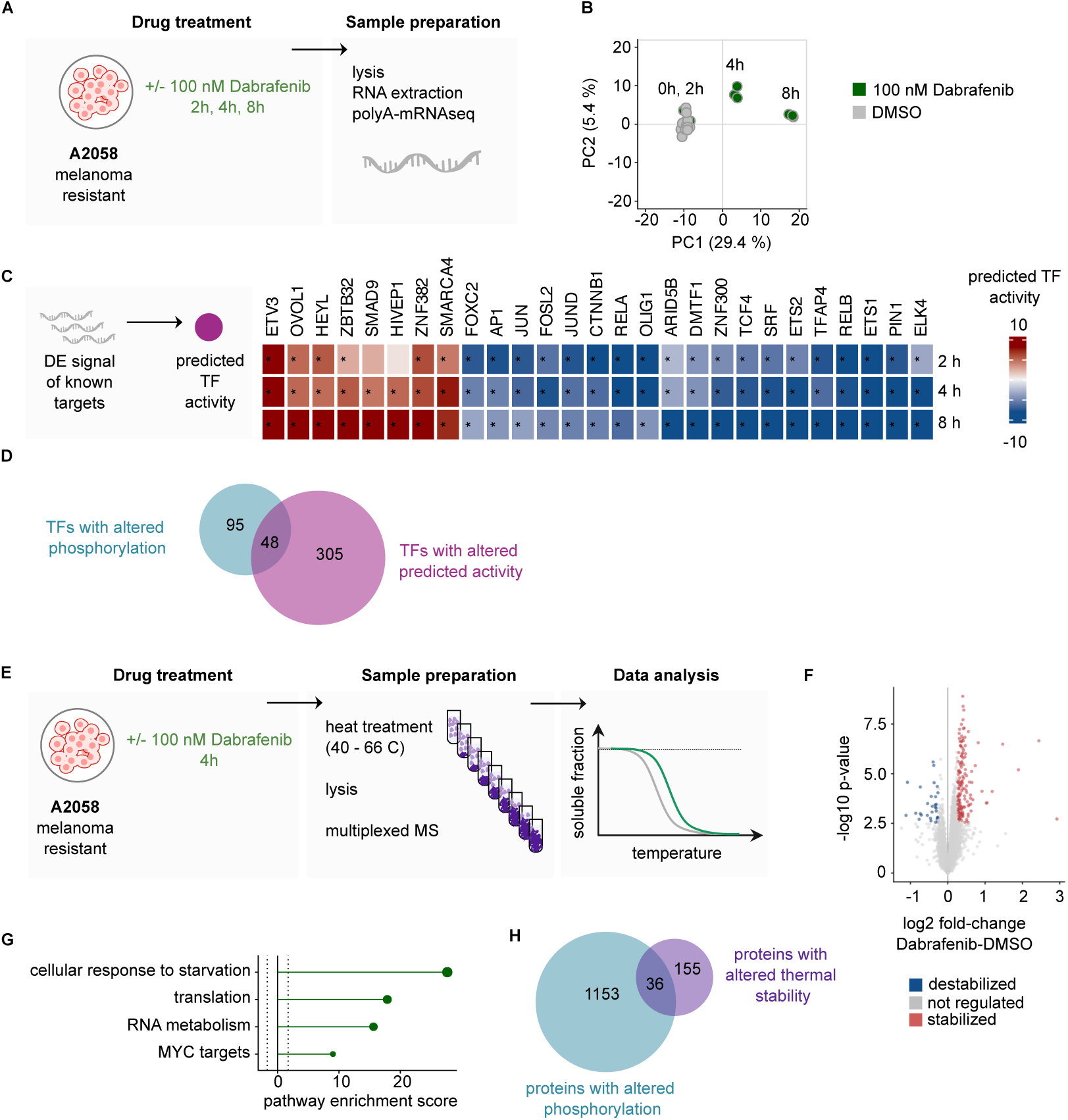
**(A)** Overview of the time-resolved transcriptomic profiling workflow. A2058, COLO-201 and MNT-1 cells (results here shown for A2058) were treated with 100 nM Dabrafenib or DMSO, and samples were collected at 0, 2, 4, and 8 hours for poly(A)-mRNA sequencing. Transcript abundance dynamics were used to infer the transcriptional programs downstream of BRAF inhibition. **(B)** PCA of gene counts of the top 10% most variable genes in Dabrafenib- (green) and DMSO-treated (grey) A2058 cells. **(C)** Inference of transcription factor (TF) activity using gene expression signatures upon Dabrafenib treatment (2-8 h) of known targets (normalized weighted mean test with CollecTRI database, p-value < 0.05). Heatmap shows scores of top TFs across time points for A2058 cells, highlighting both upregulated (red) and downregulated (blue) TFs. **(D)** Venn diagram showing the overlap between TFs with significantly altered phosphorylation in the Dabrafenib phosphoproteomics time course and TFs with inferred changes in activity upon Dabrafenib treatment based on the transcriptomics data. **(E)** Schematic overview of the thermal proteome profiling (TPP) experiment. A2058 cells treated with 100 nM Dabrafenib or DMSO for 4 hours were subjected to a temperature gradient, followed by lysis and multiplexed MS-based quantification of soluble proteins across the temperature range. **(F)** Volcano plot of differential protein thermal stability upon Dabrafenib treatment (100 nM, 4 h) relative to DMSO. Blue and red dots denote stabilizing and destabilizing responses, respectively (moderated t-test, adjusted p-value < 0.05). **(G)** Pathway enrichment analysis of proteins showing altered thermal stability upon Dabrafenib treatment (normalized weighted mean test, with GO terms and Reactome pathways, p-value < 0.05). **(H)** Venn diagram showing overlap between proteins with altered phosphorylation in the Dabrafenib phosphoproteomics time course and those with altered thermal stability in TPP upon Dabrafenib treatment.

**Figure S5.**
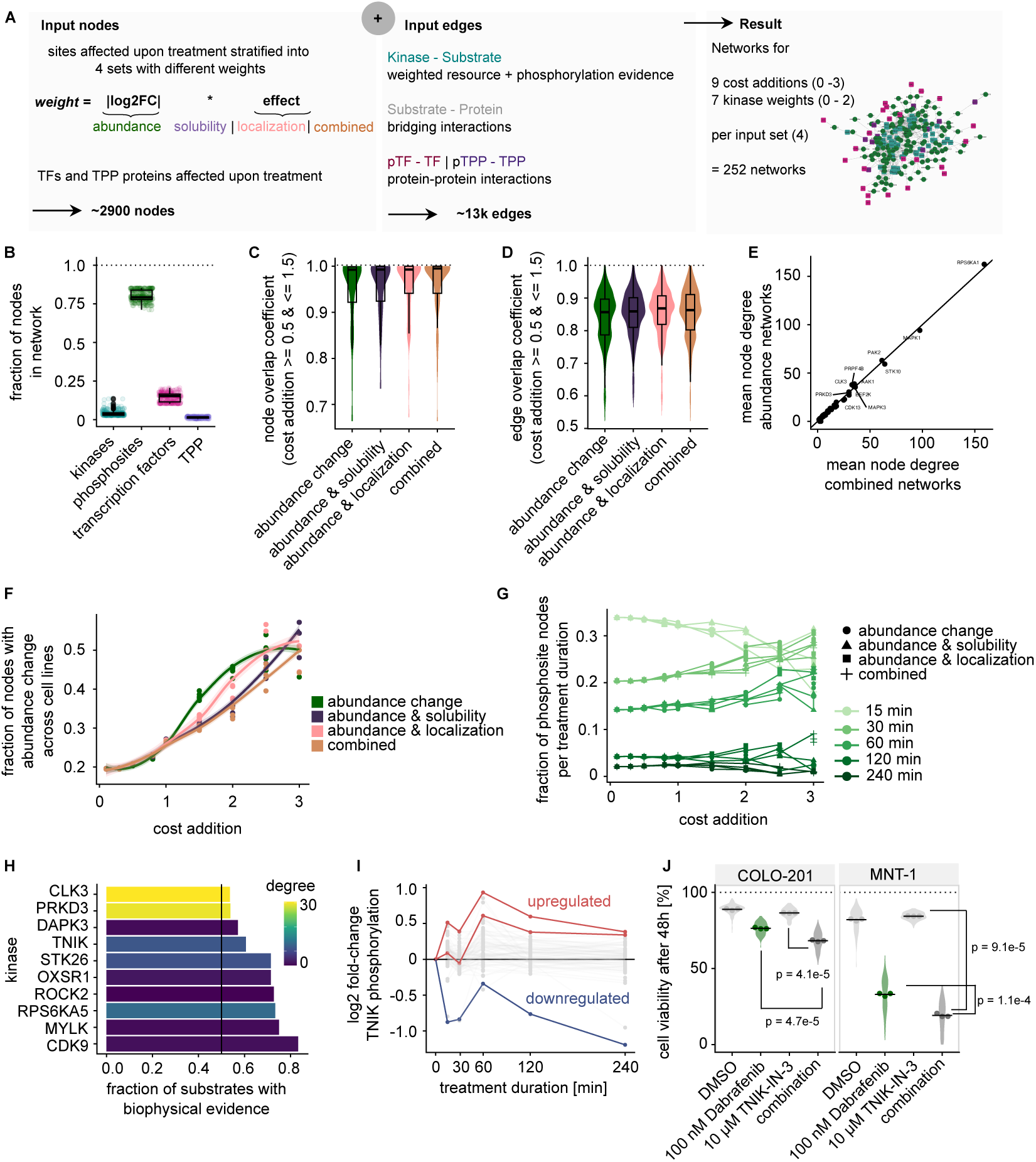
**(A)** Overview of network construction. Left: Input node preparation. Around 2500 phosphosites affected upon Dabrafenib treatment are stratified into four sets with different weights (abundance, solubility-abundance, localization-abundance, combined biophysical evidence). Transcription factors (TFs) and thermal proteome profiling (TPP) altered upon Dabrafenib treatment are included as additional nodes (∼2900 nodes total). Middle: Prior knowledge network (PKN) preparation comprising kinase-substrate interactions (weighted for measured kinase phosphorylation), protein-protein interactions, and phosphorylation evidence (∼13k edges). Right: Resulting directed multi-omic networks generated across different kinase weights and cost additions and input sets (∼252 networks total) with 1454 - 41 edges. Example network shown. **(B)** Fraction of nodes per input modality (phosphosites, TFs, TPP) or inferred kinases. Each point represents one network. Boxplots indicate mean, first and third quartiles. **(C–D)** Edge and node overlap coefficients across all networks for a given input set. Boxplots indicate mean, first and third quartiles. **(E)** Comparison of node degree centralities for networks without and with combined biophysical evidence weighting strategy. **(F)** Fraction of sites affected globally across cell lines (Figure 1B) in abundance-only, abundance-solubility, abundance-localization, or combined networks. **(G)** Fraction of nodes per input set (abundance, solubility, localization, combined) across different edge costs for sets of phosphosites reaching 20% of their maximum observed fold-change at different time points during Dabrafenib treatment. **(H)** Fraction of substrates with biophysical evidence for the top kinases in the network model coloured by overall network degree. Black line indicates 50%. **(I)** Temporal phosphorylation dynamics in the Dabrafenib time course data of TNIK coloured by direction of regulation (S570 and S678 upregulated, S795 downregulated). **(J)** Cell viability after 48 hours in COLO-201 and MNT-1 cells treated with Dabrafenib (100 nM) alone, TNIK-IN-3 alone (10 µM) or the combination (unpaired, two-sided t-test). The experiment was performed in three biological replicates (points), 56 data points were generated per replicate and condition (violin shape). Black lines indicate the mean.

**Figure S6.**
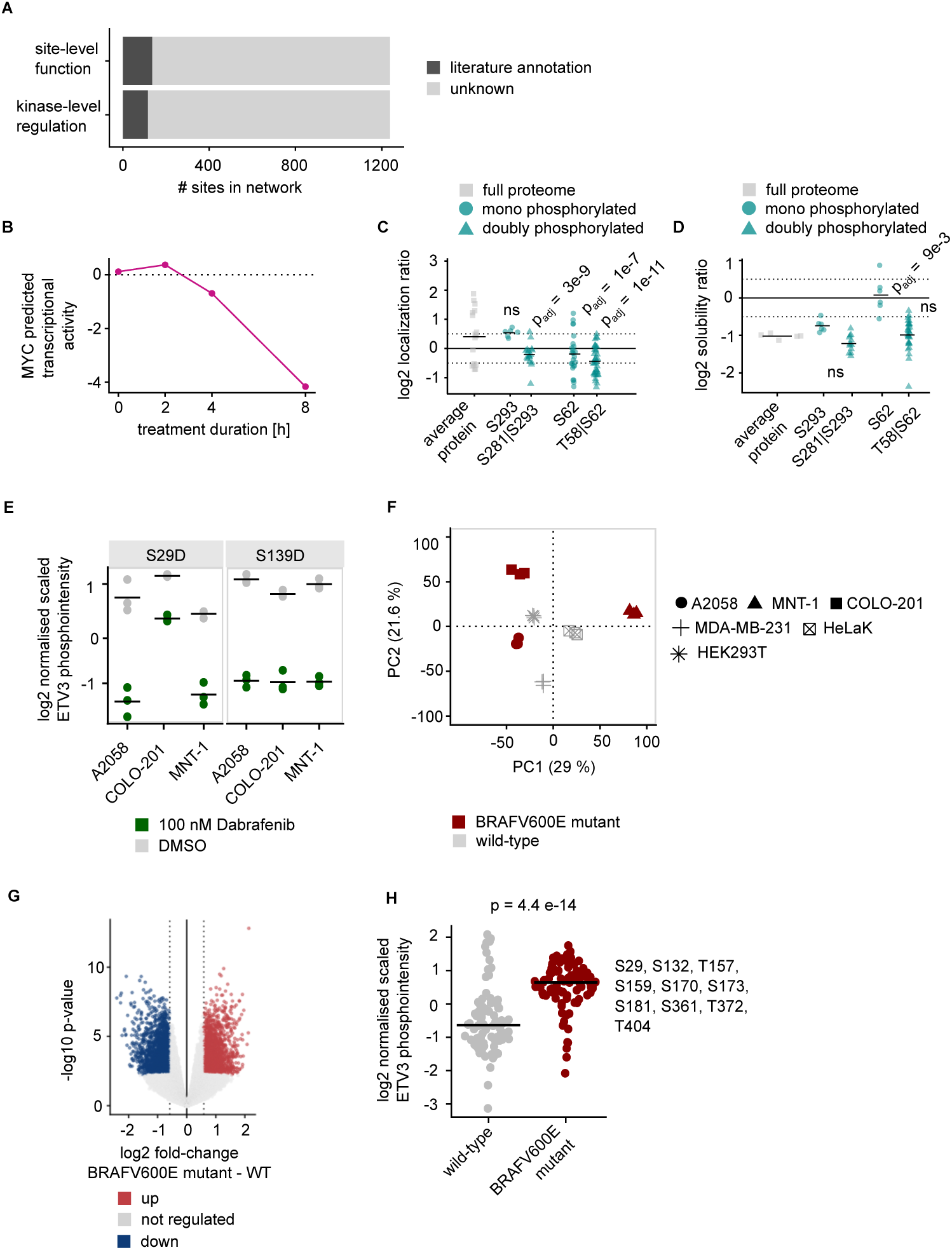
**(A)** Number of sites across all network models, coloured by whether prior annotation on site function or an upstream kinase is available in literature. **(B)** Time-resolved predicted MYC transcription factor activity following Dabrafenib treatment in A2058 cells. **(C)** log₂ localization ratios for the MYC protein and phosphopeptides in A2058 cells. Significance was assessed using limma’s moderated t-test. Black lines indicate the mean. **(D)** log₂ solubility ratios for the MYC protein and phosphopeptides in A2058 cells. Significance was assessed using limma’s moderated t-test. Black lines indicate the mean. **(E)** Normalised scaled phosphorylation levels of ETV3 S139 and S29 across BRAFV600E-mutant cell lines (A2058, MNT-1, COLO-201) treated with Dabrafenib versus DMSO. Black lines indicate the mean. ETV3 dephosphorylation is significant in all cell lines (moderated t-test, adj. p-value < 0.05, exact p-values in Supplementary Table S2). **(F)** PCA of normalized phosphopeptides of three BRAFVE600E-mutant (red) and three other cell lines (grey). The PCA was calculated based on the top 10% most variable phosphopeptides across samples. **(G)** Volcano plot of differential phosphopeptide abundance in BRAFV600E-mutant (A2058, MNT-1, COLO-201) relative to cells without the mutation (HelaK, HEK293T, MDA-MB-231). Significantly upregulated (red) and downregulated (blue) phosphopeptides are highlighted (moderated t-test, adj. p-value < 0.05, |log₂ fold-change| > log₂(1.5) as indicated by dashed line). **(H)** Expression levels of phosphorylated ETV3 in steady state across all quantified sites, replicates and cell lines for BRAFV600E-mutant cells and cells without the mutation. Black lines indicate the mean. Significance was assessed using unpaired, two-sided t-test.

**Figure S7.**
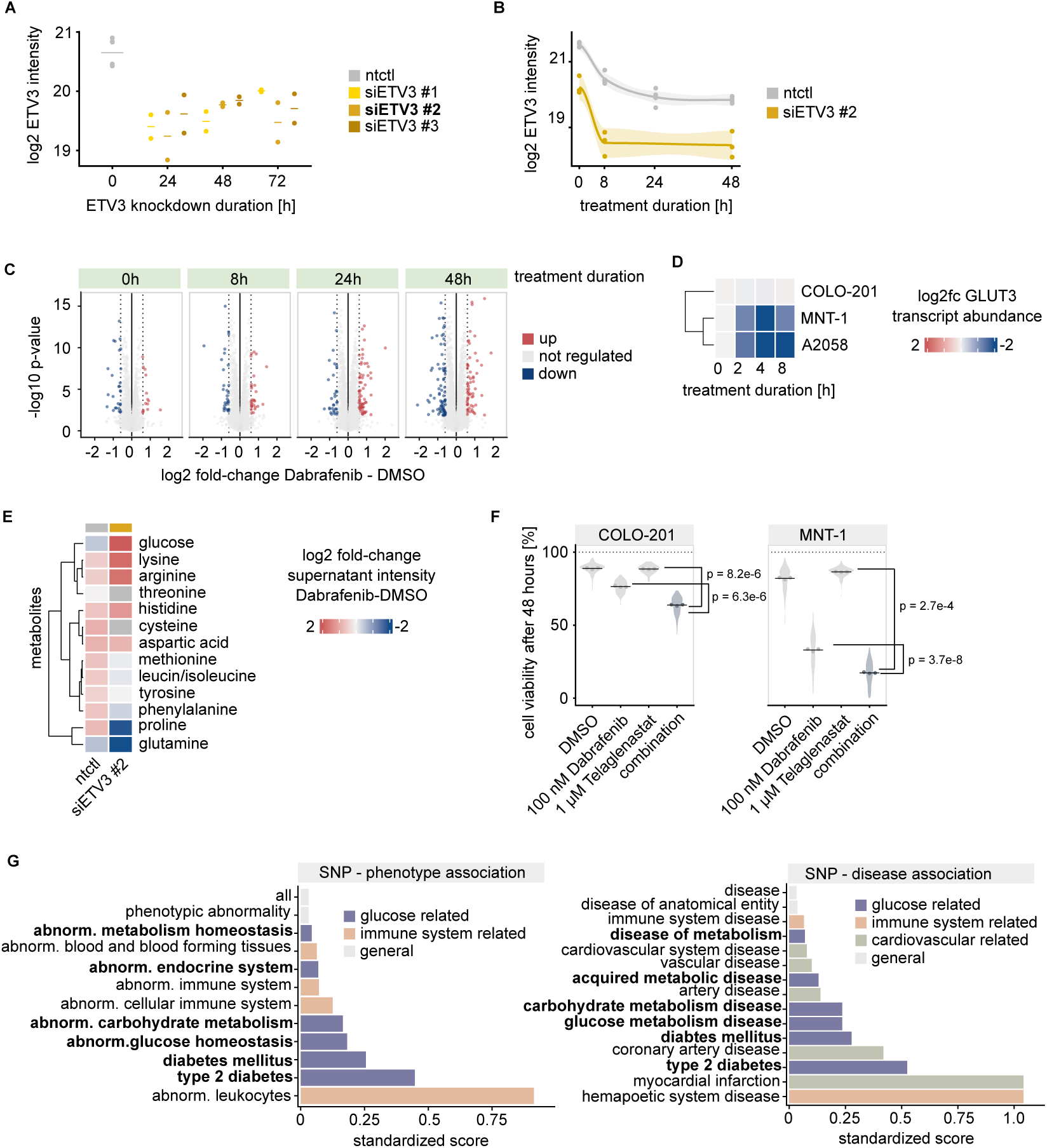
**(A)** Validation of ETV3 knockdown using three independent siRNAs. All siRNAs efficiently reduce ETV3 protein levels compared with non-targeting control (ntctl). siETV3#2 was chosen for further experiments. Central lines indicate the mean. **(B)** Temporal profile of ETV3 protein abundance upon Dabrafenib treatment and knockdown in A2058 cells. After 24 hours of treatment, ETV3 expression cannot be detected anymore. Fit lines indicate mean protein intensities (loess fit). **(C)** Volcano plots of differential protein abundance in siETV3 and control A2058 cells at 0, 8, 24, and 48 hours of Dabrafenib treatment. Significantly upregulated (red) and downregulated (blue) phosphopeptides are highlighted (moderated t-test, adj. p-value < 0.05, |log₂ fold-change| > log₂(1.5) as indicated by dashed line). **(D)** Transcript expression changes of glucose transporters GLUT1 and GLUT3 across three BRAFV600E-mutant cell lines in response to Dabrafenib. **(E)** Heatmap of quantified metabolite levels in cellular supernatant of A208 cells after Dabrafenib treatment in control versus siETV3 A2058 cells. Central lines indicate the mean. **(F)** Cell viability after 48 hours in COLO-201 and MNT-1 cells treated with Dabrafenib (100 nM) alone, Telaglenastat (1 μM) alone or the combination (unpaired, two-sided t-test). The experiment was performed in three biological replicates (points), 56 data points were generated per replicate and condition (violin shape).Central lines indicate the mean. **(G)** Standardized scores indicating significant ETV3 single-nucleotide polymorphisms (SNPs) to phenotypes or diseases associations in the GWASdb. Colours show phenotype/disease group.

## Methods

### Experimental procedures

Experiments were performed at room temperature if not stated otherwise.

#### Cell lines

All cell lines used in this study were verified to be negative for mycoplasma contamination and all cells were grown at 37 °C and 5% CO2.

A2058 cells (ATCC, CRL-11147) were cultured in Dulbecco’s Modified Eagle Medium (DMEM; Sigma-Aldrich, D5648) containing 4.5 mg mL−1 glucose and 10% (v/v) fetal bovine serum (FBS; Gibco, 10270) at 37 °C with 5% CO2. MNT-1 cells (ATCC, CRL-3450) were cultured in DMEM (Sigma-Aldrich, D5648) containing 4.5 mg mL−1 glucose, 10% (v/v) FBS (Gibco, 10270) and 1 mM L-glutamine (Gibco, 25030081) at 37 °C with 5% CO2. COLO-201 cells (ATCC, CCL-224) were cultured in Roswell Park Memorial Institute medium (RPMI; Gibco, 31800089) containing 10% (v/v) FBS (Gibco, 10270) at 37 °C with 5% CO2. These three cell lines were chosen to represent different BRAF inhibition sensitivity levels as well as different tissues of origins and phenotypes (adherence and melanin production). HeLa Kyoto cells (a kind gift by the Ellenberg Lab, EMBL), MDA-MB-231 cells (a kind gift by the Krijgsveld lab) and HEK293T (ATCC, CRL-3216) cells were cultured in DMEM (Sigma-Aldrich, D5648) containing 4.5 mg mL−1 glucose, 10% (v/v) FBS (Gibco, 10270) and 1 mM L-glutamine (Gibco, 25030081) at 37 °C with 5% CO2.

#### Drug treatments and cell viability measurements

For phosphoproteomic and transcriptomic experiments, three replicates of cells per timepoint were seeded in 150 mm dishes and grown to a density of 80%. One hour prior to treatment, the concentration of FBS was reduced to 2%. 100 mM of Dabrafenib (Selleckchem, S2807) were added at the indicated timepoints (Figure 1) in reverse order. The concentration was chosen to be in the lower range of the estimated plasma concentration of Dabrafenib to induce a physiological response. Time points were chosen to capture the immediate response to the treatment without altering protein abundance levels. Control plates were treated with an equal volume of dimethyl sulfoxide (DMSO). To stop the treatment, plates were placed on ice, washed with ice-cold Phosphate buffered saline (PBS; 2.67 mM KCl, 1.5 mM KH2PO4, 137 mM NaCl and 8.1 mM NaH2PO4, pH 7.4) twice and collected by scraping. The cells were pelleted by centrifugation at 300 g for 3 min, flash frozen in liquid N2 and stored at −80 °C.

For cell viability experiments, cells were seeded in 6-well plates and grown to 70% density. The medium was replaced with 1 mL medium containing the respective drug concentrations (100 nM Dabrafenib (Selleckchem, S2807), 1 µM Telaglenastat (Selleckchem, S7655), 10 µM TNIK-IN-3 (HY-145293), 100 nM NVP-2 (HY-12214A), 1 µM CLK3-IN-T3 (HY-115470) or an equal volume of DMSO as control) and 1 µL of 5 mM SYTOX green (Invitrogen™, S7020). Cells were treated for up to 48 hours, To assess cell viability, the fluorescence-based and lysis-dependent inference of cell death kinetics (FLICK) protocol ^65^ was applied using a BioTek Synergy H1 plate reader. SYTOX intensity was measured with a 70% gain at 72 positions per well. After 48 hours of measurement/treatment, cells were lysed and the total SYTOX intensity was determined to calculate the total cell signal.

#### siRNA transfection experiments

ETV3 small inhibitory RNAs (siRNAs) and a non-targeting control siRNA used were acquired from ThermoFisher™. Three siRNAs were tested and one was chosen for the time course experiment (Supplementary Table 8). Cells were plated one day before siRNA transfections, targeting 50% confluency on the day of treatment. The Lipofectamine RNAiMAX protocol (Thermo Fisher Scientific, 13778075) was employed following the manufacturer’s manual. For a single 6-well plate reaction, 150 μL of Opti-MEM™ (Gibco, 31985070) were mixed with 6 μL of Lipofectamine RNAiMAX reagent. Separately, 20 μmol siRNA were diluted with 150 μL of Opti-MEM™. The mixtures were combined to a transfection mixture and incubated for 15 min at room temperature (RT). Cell medium was replaced by 700 µL Opti-MEM™ and 300 µL of the transfection mixture was dripped into each well. After 24 hours, the medium was replaced with standard medium. For combined knockdown and Dabrafenib treatment experiments, the medium was replaced with standard medium containing 100 nM Dabrafenib or an equal volume of DMSO after 48 h. For proteomics experiments, cells were treated for the indicated durations (Figure 6), washed with PBS (2.67 mM KCl, 1.5 mM KH2PO4, 137 mM NaCl and 8.1 mM NaH2PO4, pH 7.4) twice and collected by scraping. The cells were pelleted by centrifugation at 300 g for 3 min. The cell pellets were flash frozen in liquid N2 and stored at −80 °C.

For metabolomics experiments, cells were treated with 100 nM Dabrafenib or an equal volume of DMSO for 24 hours after 48 hours of knockdown. The supernatant was transferred to 1.5 mL tubes and centrifuged at 300 g for 3 min to remove cell debris. 500 µL supernatant per sample was flash frozen in liquid N2 and stored at −80 °C. The cells were harvested by trypsinization and centrifugation at 300 g for 3 min and lysed in 2% Sodium dodecyl sulfate (SDS) for 10 min at 96 °C. Protein concentrations were determined using the Pierce™ Dilution-Free™ Rapid Gold BCA Protein Assay (Thermo Scientifc, A55860).

#### (Phospho)proteomics sample preparation

Frozen cell pellets were lysed in a buffer constituted of 4 M guanidinium isothiocyanate (Gnd-SCN), 50 mM 2-[4-(2-hydroxyethyl)piperazin-1-yl]ethanesulfonic acid (HEPES), 10 mM tris(2-carboxyethyl)phosphine (TCEP), 1% N-lauroylsarcosine, 5% isoamyl alcohol and 40% acetonitrile (ACN) adjusted to pH 8.5 with 10 M NaOH. The volume of the used lysis buffer corresponded to approximately 5 times the sample volume. After 1 hours of incubation in a shaker at 1,000 rpm, samples were centrifuged at 16,100 g for 10 min or filtered in multiscreenHTS-HV 0.45 µm 96-well filter plates with polyvinylidene fluoride (PVDF) membranes (Merck Millipore) to remove cell debris and nucleic acid aggregates. Protein concentrations were determined using tryptophan fluorescence assay^66^ and the samples were diluted to a maximum concentration of 25 µg/µL, if needed. Volumes of 80 µL of lysate per well were transferred into a multiscreenHTS-HV 0.45 µm 96-well filter plate, and 220 µL of ice-cold ACN were added to induce protein precipitation. After 10 min, the plate was centrifuged at 1,000 g for 2 min to remove solution and protein precipitates, were washed and centrifuged twice with 200 µL 80% ACN and two times with 200 µL 70% EtOH. Then, the digestion buffer composed of 50 mM Triethylammonium bicarbonate buffer (TEAB) and TPCK treated trypsin (Thermo Fisher Scientific, 20233) was added to the protein precipitates in the filter plate. The ratio of trypsin to protein was fixed to 1:25 w/w and the maximum final protein concentration to 10 µg/µL. Trypsin was reconstituted at a stock concentration of 10 µg/µL and added to the ice-cold digestion buffer shortly prior to the addition to the precipitates. Tryptic digestion was carried out under mild shaking at 600 rpm overnight.

For the phosphoproteomics Dabrafenib time course, the Dabrafenib treatment of three BRAFV600E cell lines and the steady state cell line profiles, triplicates of the experiment were multiplexed into one Tandem mass tag pro 18-plex (TMTpro 18) set for the full proteome and the phosphoproteome.

#### Solubility proteome profiling

Lysis buffer was composed of PBS containing 1 U mL−1 RNase inhibitor (RNasin Plus, N2615), cOmplete protease inhibitor (Roche), PhosSTOP phosphatase inhibitor (Roche), 1.5 mM MgCl2 and 0.8% NP-40. Frozen A2058 cell pellets were thawed on ice and resuspended in lysis buffer twice the volume of the pellet. This homogeneous cell suspension was subjected to mechanical disruption by three cycles of freezing in liquid N2 and thawing at 25 °C. Protein concentrations were determined using tryptophan fluorescence assay. The lysates were diluted to the lowest sample concentration measured and each sample was divided into one 110 µL and one 90 µL aliquot. The total proteome aliquot of 90 µL was solubilized with SDS at a final concentration of 1%. The soluble proteome aliquots of 110 µL were centrifuged at 100,000 g for 20 min at 4 °C. 90 µL of supernatant containing the soluble pool of proteins were obtained.

ACN was added to all samples to a final concentration of 70% to induce protein precipitation. Supernatant was removed and precipitates incubated in 80 µL of a buffer constituted of 4 M guanidinium isothiocyanate, 50 mM HEPES, 10 mM TCEP, 1% N-lauroylsarcosine, 5% isoamyl alcohol and 40% ACN adjusted to pH 8.5 with 10 M NaOH under shaking conditions at 100 rpm for one hour at RT. 80 µL of lysate per well were transferred into a multiscreenHTS-HV 0.45 µm 96-well filter plate, and 220 µL of ice-cold ACN were added to induce protein precipitation. The samples were then washed and digested as previously described in section (phospho)proteomic sample preparation. Triplicates of the experiment were multiplexed into one TMTpro 18-plex for the full proteome and the phosphoproteome.

For the SPP experiments combined with a DNA digestion step, the soluble proteome aliquot was split into two and to one 1 uL of DNAse 1 (Qiagen, 79254) was added for 1 hour at 4 °C. The samples were afterwards processed as described above.

#### Nuclear enrichment profiling

Lysis buffer was composed of PBS containing cOmplete protease inhibitors (Roche), PhosSTOP phosphatase inhibitors (Roche) and 0.8% NP-40. Fresh A2058 cell pellets were resuspended in a volume of lysis buffer equal to twice the volume of the pellet by pipetting ten times. This homogeneous cell suspension was incubated ice for one minute before centrifugation at 4000 g for 30 s at 4 °C. The pellet corresponding to the nuclear enriched fraction was separated from the supernatant immediately. The supernatant samples were solubilized with SDS with a final concentration of 1%, and ACN was added to a final concentration of 70% to induce protein precipitation. Both the nuclear enriched pellet and the supernatant precipitates were incubated in 200 µL of a buffer constituted of 4 M guanidinium isothiocyanate, 50 mM HEPES, 10 mM TCEP, 1% N-lauroylsarcosine, 5% isoamyl alcohol and 40% ACN adjusted to pH 8.5 with 10 M NaOH for one hour on a shaker at RT at 100 rpm.

Protein concentrations were determined using tryptophan fluorescence assay. Volumes of 80 µL of lysate per well, corresponding to 500 ng nuclear enriched and 200 ng supernatant sample, were transferred into a multiscreenHTS-HV 0.45 µm 96 well filter plate, and 220 µL of ice-cold ACN were added to induce protein precipitation. The samples were then washed and digested as described in section (phospho)proteomic sample preparation. Triplicates of the experiment were multiplexed into one TMTpro 18-plex for the full proteome and the phosphoproteome.

Please note, that nucleoplasmic proteins not tightly bound to chromatin can leak into the cytoplasmic fraction during extraction, potentially confounding nuclear enrichment data (e.g. observed for *ETV3*).

#### Thermal proteome profiling

Thermal proteome profiling (TPP) was performed in live A2058 cells in a TPP-TR setting as described before67. In brief, cells were treated with 100 nM Dabrafenib or DMSO and incubated at 37 °C and 5% CO2 for 4 hours, then harvested by trypsinization and centrifugation. Cells were resuspended in PBS and transferred to 96-well PCR plates. Cells were heated for 3 min at one of the 9 tested temperatures (40.4, 44.0, 46.9, 49.8, 52.9, 55.5, 58.6, 62.0 and 66.3 °C) and incubated for 3 min at RT thereafter. Cells were lysed with PBS containing cOmplete protease inhibitor (Roche), PhosSTOP phosphatase inhibitor (Roche), 0.8% NP-40, 2 mM MgCl2 and benzonase 1 kU ml−1. After a 30 min incubation on ice, the plate was centrifuged at 3,000 g for 10 min at 4 °C and the supernatant was transferred to a multiscreenHTS-HV 0.45 µm 96-well filter plate with PVDF membranes (Merck Millipore). After filtration, the samples were denatured by addition of the same volume of denaturing buffer (6 M Gnd-SCN, 100 mM HEPES, 10 mM TCEP) and the protein precipitated by addition of ACN to a final concentration of 70%. The samples were then washed and digested as previously described in the sample preparation section. Protein concentrations were determined using tryptophan fluorescence assay. 10 µg of protein per sample were used for TMT labeling. Dabrafenib treated samples were labeled with channels 1-9 while DMSO treated samples were labeled with channels 10-18 before pooling. Three biological replicates were prepared.

#### Phosphopeptide enrichment

For phosphoproteomics studies, including the time course experiments, experiments investigating different cell lines in steady state and in response to Dabrafenib, phospho-SPP experiments and nuclear fractionation, phosphopeptide enrichment was performed. Lyophilized peptides were resuspended in a loading and washing buffer composed of 80% ACN and 0.07% trifluoroacetic acid (TFA), sonicated, and centrifuged at 1000 x g for 30 seconds. Phosphopeptide enrichment was carried out using the supernatant as described by Bartolec at al ^99^ using the KingFisher Apex robot (Thermo Fisher Scientific) with 25 µL of Fe-NTA MagBeads (PureCube) per sample. Phosphopeptides were eluted with 100 µL of 0.2% diethylamine in 50% ACN, followed by lyophilization. The enriched phosphoproteomic samples were TMT labelled see section TMT labeling) and then subjected to offline Porous graphitic carbon based Liquid Chromatography (PGC-LC).

#### TMT Labeling

For the full proteome measurements, 10 µg of peptides per condition were labeled with TMT, while for the phosphoproteome, the totality of enriched phosphopeptides was used. Peptides were resuspended in 10 µL of 100 mM HEPES (pH 8.5), and 2 µL (phosphopeptides) or 4 µL (full proteome) of TMTpro 18-plex reagent at a concentration of 20 µg/µL in ACN were added. The labeling reaction was allowed to proceed for 1 hours at RT, and then was quenched by the addition of 5 µL of 5% hydroxylamine for 15 min. Labeled peptides belonging to the same experiment were subsequently pooled and lyophilized. Prior to fractionation, the phosphopeptides were resuspended in 50 µL of 10% TFA and desalted using in-house C18 stage tips packed with 1 mg of ReproSil-Pur 120 C18-AQ 5 µm material (Dr. Maisch) above a C18 resin plug (AttractSPE disks bio - C18, Affinisep), while in the case of the full proteome samples, the peptides were desalted on an OASIS plate (Waters).

#### PGC-LC offline peptide fractionation

The samples were resuspended in 18 µL of buffer A (0.05% TFA in MS-grade H2O supplemented with 2% ACN), with an injection volume set to 16 µL. To separate peptides, PGC fractionation was performed using a Hypercarb column (100 mm length, 1.0 mm inner diameter, 3 µm particle size, Thermo Fisher Scientific) at 50 °C, with a flow rate of 75 µL/min, on an Ultimate 3000 LC system (Thermo Fisher Scientific). After a 1 min post-injection delay, a linear gradient separation was applied, increasing from 13% buffer B (0.05% TFA in ACN) to 42% buffer B over 47 and 95 min for the phosphoproteome and full proteome samples, respectively. This was followed by an increase to 80% buffer B within 5 min. The column was then washed with 80% buffer B for 5 min before re-equilibration with 100% buffer A for another 5 min. Fractions were collected from 4.5 to 52.5 min and from 4.5 to 100.5 min at 2 min intervals, yielding 24 fractions and 48 fractions for the phosphoproteome and full proteome samples, respectively. These were then pooled into 24 fractions by combining each fraction with its corresponding n + 24 fraction for the phosphoproteome and full proteome samples, respectively. After pooling, the fractions were lyophilized using a vacuum concentrator prior to liquid chromatography coupled to tandem mass spectrometry (LC-MS/MS) analysis.

#### Proteomics MS

Prior to injection, all samples were resuspended in a loading buffer composed of 1% TFA, 50 mM citric acid, and 2% ACN in MS-grade water. LC separation was performed on an UltiMate 3000 RSLCnano system (Thermo Fisher Scientific). Peptides were initially trapped on a cartridge (Precolumn: C18 PepMap 100, 5 μm, 300 μm i.d. × 5 mm, 100 Å) before being separated on an analytical column (Waters nanoEase HSS C18 T3, 75 μm × 25 cm, 1.8 μm, 100 Å). Solvent A was composed of 0.1% formic acid with 3% DMSO in MS-grade water, while solvent B contained 0.1% formic acid with 3% DMSO in LC–MS-grade ACN. Peptides were loaded onto the trapping cartridge at 30 μL/min with solvent A for 5 min, then eluted at a constant flow rate of 300 nL/min using a linear gradient of buffer B, followed by an increase to 40% buffer B, a wash at 80% buffer B for 4 min, and re-equilibration to initial conditions. The linear gradient corresponded to an increase from 7% to 27% B. The LC system was coupled to either a Fusion Lumos Tribrid, an Exploris 480 mass or a Q-Exactive Plus mass spectrometer (Thermo Fisher Scientific), operated in positive ion mode. The mass spectrometers were operated in data-dependent acquisition mode with a maximum duty cycle time of 3 sec or a top 20 method, selecting precursors with charge states 2–7 and a minimum intensity of 2 × 10⁵ for subsequent Higher-Energy Collisional Dissociation (HCD) fragmentation. Peptide isolation was performed using the quadrupole with a 0.7 m/z isolation window. MS/MS spectra were acquired in profile mode using an Orbitrap mass spectrometer, with a maximum injection time of 100 ms and an AGC target of 1 × 10⁵ charges.

For the data-independent acquisition (DIA) samples (proteomics time course of combined ETV3 knockdown and Dabrafenib treatment), peptides were separated using an Vanquish Neo Ultra High-Performance Liquid Chromatography (UHPLC) system (Thermo Fisher Scientific) operated in trap-and-elute mode. The LC was equipped with a trapping cartridge (Precolumn; PepMap Neo C18, 5 μm, 300-μm inner diameter × 5 mm, 100 Å) and an analytical column (Ionopticks AUR3-25075C18-XT, 25 cm x 75 μm ID, 1.7 μm C18). Solvent A consisted of 0.1% formic acid supplemented with 3% DMSO in LC–MS-grade water and solvent B was 0.1% formic acid supplemented with 3% DMSO in LC–MS-grade ACN. Peptides were loaded onto the trapping cartridge and separated on the analytical column using a linear gradient of 5–26% of buffer B at a flow rate of 300 nL/min for 29.7 min, followed by an increase to 40% of buffer B within 3 min before washing at 85% with buffer B for 5 min and re-equilibration to initial conditions, resulting in a total MS acquisition time of 40 min. The LC system was coupled to an Orbitrap Astral mass spectrometer (Thermo Fisher Scientific) operated in data independent acquisition mode. The instrument was operated in positive ion mode with a spray voltage of 1.8 kV and a capillary temperature of 280 °C. Full-scan MS spectra were acquired in the Orbitrap at a resolution of 240,000 with a mass range of 430-680 m/z, an automatic gain control (AGC) target of 5e6 charges and a maximum injection time of 5 ms. DIA spectra were acquired in the Astral mass analyzer with 2 m/z windows between 430 and 680 m/z. MS2 scan range was set to 150-2000 m/z, the normalized collision energy to 25 and the default charge state to 2+. The normalized AGC target was set to 500% and the maximum injection time to 5 ms.

#### Metabolomics sample preparation

Samples were prepared for LC-MS analysis by organic solvent extraction. In brief, 20 μL of supernatant was mixed with 105 µL of organic solvent (acetonitrile:methanol, 1:1) supplemented with an internal standard mixture. Internal standard mixture consisted of phenylalanine-d5, tryptophan-d5, ibuprofen-d4, tolfenamic acid-d4, estriol-d3, diclofenac-d4, warfarin-d5, oxfendazole-d3, chloramphenicol-d5, nafcillin-d5 and caffeine-d9, each to a final concentration of 80 nM. The material was homogenized and incubated at -20°C to allow protein precipitation. After incubation, samples were centrifuged (3220 g, 4°C) for 15 min. 15 µL of supernatant was diluted with 30 µL of the starting condition of the respective HILIC conditions (see below).

#### Metabolomics LC-MS acquisition

Liquid chromatography-mass spectrometry analysis was carried out as previously described ^68^. Normal-phase chromatographic separation was acquired using an InfinityLab Poroshell 120 HILIC-Z, 2.1 x 150 mm, 2.7-micron pore size column (Agilent Technologies, U.S.A.), in which different separation strategies and MS parameters were used for positive and negative ionization modes. For positive MS, the mobile phases were (a) water with 5 mM ammonium formate and 0.1% (v/v) formic acid, and (b) acetonitrile, 5 mM ammonium formate, and 0.1% (v/v) formic acid as solvent B. The column compartment was kept at 25 °C. 5 µL sample were injected at 98% B and 0.250 ml/min flow, followed by 98% B until minute 3, a gradient to 70% B until minute 11, a gradient to 60% B until minute 12, a gradient to 5% B to minute 16, and remaining at 5% B until minute 18 before re-equilibration to 98% until minute 20. The qTOF (Agilent) was operated in positive scanning mode (50-1500 *m/z*) with the following source parameters: VCap = 3000 V; nozzle voltage = 0 V; gas temp = 225 °C; drying gas flow = 11 l/min; nebulizer = 40 psig; sheath gas temp = 225 °C; sheath gas flow = 10 l/min; fragmentor = 300 V; and Octopole RF Vpp = 450 V. Online mass calibration was performed using a second ionization source and a constant flow of reference solution (reference ions are *m/z* 121.0508 and 922.0097, Supplementary Table 7).

#### Transcriptomics

Approximately 8*10^6 cells were lysed in 600 µL RLT buffer (Qiagen, #74181) and RNA was extracted using the RNeasy 96-well plate kit (Qiagen, #74181) according to manufacturer’s instructions including the DNase step.

The initial RNA was QCed using Agilent Bioanalyzer with the RNA Nano Assay kit as per the manufacturer’s protocol^69^. The RNA sample set was then standardized to 200 ng total RNA in 50 µL using the concentration values given by the Bioanalyzer. The libraries were prepared on a Beckman Coulter Automated Workstation Biomek i7 Hybrid (MC + Span-8). For library preparation an automated version of the NEBNext® Ultra™ II Directional RNA Library Prep Kit was used, following section 1 - Protocol for use with NEBNext Poly(A) mRNA Magnetic Isolation Module ^70^. An adaptor dilution of 1 to 30 was used, the samples were individually barcoded using unique dual indices during the PCR using 12 PCR cycles as per the manufacturer’s protocol. The individual libraries were quantified using the Qubit HS DNA assay as per the manufacturer’s protocol. For the measurement 1 µL of sample in 199 µL of Qubit working solution was used. The quality and molarity of the libraries was assessed using Agilent Bioanalyzer with the DNA HS Assay kit as per the manufacturer’s protocol. The assessed molarity was used to equimolarly combine the individual libraries into one pool for sequencing. The pool was loaded and sequenced on an Illumina NextSeq 2000 platform (Illumina, San Diego, CA, USA) using a P3 100 cycle kit, a read-length of 122bp single-end reads and 650pM final loading concentration.

### Data analysis

#### Cell viability

Raw SYTOX intensities were exported for analysis (45 measurements per well). For final cell number determination, outlier values falling outside the measurement range were set to the maximum detected plate intensity. Cell viability was then calculated as *1 – (measured SYTOX intensity at a given treatment duration / total SYTOX signal at endpoint)*. Values below zero were set to zero. Ratios were normalized by median normalization and a mean per well was calculated. Statistical significance between conditions was assessed using a two-sided unpaired t-test on the means of three biological replicates.

#### Database search

For all proteomic analyses, the Swissprot *Homo sapiens* database (20,443 entries) was used for the database searches. For plasmid transfection experiments, sequences of proteins encoded on the plasmid were added.

Files were searched using MSFragger ^71^ v4.0 in Fragpipe v21. The default MSFragger phospho or TMT16-phospho workflow was used, with a few modifications: oxidation on methionine (maximum 2 occurrences), phosphorylation on S/T/Y (maximum 3 occurrences) and peptide n-terminal TMT16 labeling (maximum 1 occurrence) were set as variable modifications, with a total of up to 5 variable modifications allowed per peptide. Lysine TMT16 labeling and cysteine carbamidomethylation were set as fixed modifications. Percolator ^72^ was used for PSM validation and PTMProphet ^72^ was used to determine site localization. An FDR cutoff of 1% was used. For the TMT quantification, Philosopher ^73^ was used to extract MS1 and TMT intensities.

DIA Astral raw files were analyzed using DIA-NN 19.1 with directDIA module using an in silico DIA-NN predicted spectral library (C carbamidomethylation and N-terminal M excision; 2 missed cleavage; precursor m/z range: 430-680). The spectral library was generated from the fasta file mentioned above. The DIA-NN search used the following parameters: Precursor FDR (%)= 1; Mass accuracy, MS1 accuracy, and Scan window = 0; Use isotopologues; MBR enabled; Heuristic protein inference enabled; No shared spectra; Protein inference = Genes, Neural network classifier: Single-pass mode; Quantification strategy: Robust LC (high precision); Cross-run normalization: RT-dependent; Library generation: Smart profiling; Speed and RAM usage: optimal results.

#### (Phospho)proteomics data analysis

For TMT analyses, the PSM tables produced by FragPipe were filtered to keep PSMs having full quantification (signal in all utilised TMT channels). For all experiments, contaminant proteins were filtered out. The summarisation of quantitative information at the (phospho-)peptide levels was performed by summing the TMT intensities of all the PSMs assigned to that specific (phospho-)peptide. We do report the highest localization score for each site across redundant peptide spectrum matches (PSMs), but keep all phosphopeptides in the downstream analysis for hypothesis generation. For site-level follow-up experiments, localization scores have to be taken into account and are provided in Supplementary Table 9.

For the phospho-SPP data, solubility ratios (ratio = soluble intensity/total intensity) were calculated for downstream analysis and visualisation. Outlier values for proteins and phosphopeptides were removed if their median solubility exceeded 2 or if their solubility ratio was more than two standard deviations from the median solubility. For the quantitative proteomics and phosphoproteomics data of different cell lines in steady state, we first removed the low abundant signals of phosphopeptides and proteins to avoid detection due to TMT channel leaking.

Differential expression analysis was performed using the filtered and summed TMT reporter ion intensities (summarised at the Modified Peptide level) for phosphoproteomics or using the protein intensities from MSFragger output for the full proteome experiments. As the full proteome was hardly altered with the chosen stimulation durations, phosphopeptide intensities were not corrected for protein abundance changes. Median or vsn-normalization^74^ and differential expression analysis were used as implemented in the limma ^75^ package (version 3.58.1) in R (version 4.3.3). The phospho-SPP and phospholocalization datasets were normalized per biophysical group.

For the time course experiment, limma was used to compare phosphopeptide intensities of samples to the 0 hour DMSO treated cells. For the comparison of the steady-state phosphoproteome of six cell lines, limma was used to compare protein and phosphopeptide abundance dependent on the BRAF mutation state. To assess the Dabrafenib response across three different cell lines, limma was used to compare phosphopeptide intensities of Dabrafenib treated cells to DMSO treated cells. For the phospho-SPP and phospho-nuclear fractionation datasets, intensity was modelled as the interaction between dataset (phosphopeptide or full proteome) and the sample type (solube vs total or nuclear enriched vs rest), blocking the treatment effect as there was no impact of the treatment observed. For the TPP dataset, protein intensities of Dabrafenib and DMSO treated cells were compared for a range of middle temperatures (49.8 °C, 52.9 °C, 55.5 °C) using temperature as a block factor. For experiments with ETV3 siRNAs, protein intensities of transfected cells were compared to cells transfected with a non-targeting control siRNA per tested siRNA or per Dabrafenib treatment duration. The linear models were fitted using limma’s lmFit() function and a moderated t-statistic was computed using limma’s eBayes() function. P-values were adjusted for multiple testing using the Benjamini-Hochberg (BH) procedure implemented in limma. Proteins or phosphopeptides were considered to be statistically significantly changing if they had adjusted p-values <0.05 and absolute log₂ fold-change >log₂(1.5).

#### Metabolomics data analysis

The area under the curve of glucose and glutamine was integrated using the MassHunter Quantitative Analysis Software (Agilent, version 7.0) based on the accurate high-resolution mass and RT of the reference analytes and the following parameters: signal threshold of 30,000; mass tolerances of 0.002 amu or 20 ppm; retention time tolerance of 0.3 min, for HILIC. Lastly, all metabolites annotated in this manuscript were assigned a Level 1 annotation according to the Metabolomics Standards Initiative (MSI) classification system. This includes a match in retention time using the same chromatographic system, accurate mass, and MS/MS fragmentation pattern, ensuring unequivocal identification of the compounds. Metabolite intensities were scaled with the measured protein concentration of the cell pellet to normalize for cell number differences. Scaled intensities were normalized across samples using median normalization. Normalized intensities were used to calculate ratios to the median DMSO intensity per sample. Statistical differences between ETV3 knockdown and control samples were calculated using a two-sided unpaired t-test.

#### Transcriptomic data analysis

Raw FastQ files were processed to count matrices using the nf-core nextflow pipeline for RNAseq data ^76^ (https://github.com/nf-core/rnaseq). Default parameters and the reference human genome (GRCh38.96) were used. Counts were extracted after pseudoalignment with salmon. The transcriptomics dataset was preprocessed using the DEseq2 ^77^ and RNAseqQC ^78^ packages. Genes were filtered for at least 5 counts and presence across at least 12 samples. Size factors were estimated with estimateSizeFactors(), and a design formula was specified to model the experimental treatment and control samples at each time point.

Differential expression was performed using DESeq (parallelized). For each pre-defined contrast, shrunken log₂ fold changes were obtained with lfcShrink(type = "ashr"). Genes were classified as significantly changed if |log₂ fold-change| ≥ log₂(1.5) (shrunken fold-change) and adjusted p-value < 0.05 (Benjamini-Hochberg adjusted).

#### Pathway enrichment and enzyme activity analyses

For the pathway enrichment analyses, MSIGDB Hallmark, Reactome and GO term gene sets were used with the decoupleR package ^79^ (normalized weighted mean method) to calculate pathway enrichment scores from indicated input values (for example log₂ fold-changes). This algorithm estimates an enrichment score which corresponds to the mean signal or an alternative summary statistic of all annotated pathway members. In order to estimate the activity of transcription factors (TFs) and kinases the normalized weighted mean method of the decoupleR package was used. For transcription factors, the COLLECTRI database^80^ was used to obtain TF-target interactions, downloaded using the OmnipathR package ^81^. Phosphosite-enzyme collections were assembled as described in the section kinase prior knowledge analysis. For the decoupleR tool, log₂ fold-change values or t-values with and without additional weighting of transcripts and phosphosites after DE analysis were used as input per condition. An enzyme was considered to be significantly affected for p-value < 0.05. Top regulators were used based on the absolute normalized enrichment score.

#### Structural and functional phosphosites analysis

We used the structuremap python package to generate all site-level structural annotations (v0.0.9) (https://github.com/MannLabs/structuremap) ^31^. We retrieved predicted structures for all phosphoproteins identified from the AlphaFold Protein Structure Database. Residues were annotated using predicted secondary structures and a prediction-aware part-sphere exposure (pPSE) metric described by Bludau et al, which scores the order for each residue based on the number of residues in their structural proximity, defined by a directional partial sphere with a specific radius (Å) and angle (°), which also considers AlphaFold prediction error. Two pPSE metrics were generated: One for a wider and bigger partial sphere (24Å, 180°) and other for a narrower and smaller partial sphere (12Å, 70°). As suggested, we used a threshold of 34.27 on the smoothed pPSE (180°, 24Å) to determine intrinsically disordered regions. Functional scores per phosphosite across the human phosphoproteome were predicted as described before using the funscoR R-package^30^.

#### Kinase prior knowledge analysis

Literature-based interactions were sourced from the OmniPath Enzyme-Substrate database^82^, including all phosphorylation-related interactions from PhosphoSitePlus ^64^ and SIGNOR^83^. Reported functions of phosphosites were also downloaded from PhosphositePlus. Specificity scores were gathered from the Serine/Threonine kinase library (Supplementary table 3 in Johnson et al. ^33^) and the Tyrosine kinase library (Supplementary table 3 in Yaron-Barir et al. ^32^). For computational predictions, we used Phosformer ^34^, downloading and running the model via the Python module (https://github.com/esbgkannan/phosformer). All human phosphosites listed in PhosphoSitePlus and their corresponding 11-mer sequences, centered on the phosphorylation site with six flanking amino acids on either side, were used as potential targets.

#### Network modeling and analysis

To generate the networks, we first assembled the prior knowledge network together with different sets of input nodes. For phosphosites, we considered all sites significantly regulated in the Dabrafenib time course. For the first input set, the maximum measured fold change per site was used as the prize value. In the second and third sets, this prize was doubled for sites with significant effects on protein solubility or nuclear enrichment, respectively. For the fourth set, the weighting schemes of sets 2 and 3 were combined. In all cases, transcription factors with significantly altered activity at 4 hours or 8 hours, as well as proteins with altered thermal stability, were included as additional downstream input. Finally, all scores were converted to absolute values and quantile normalized. Only kinases that showed significant changes in phosphorylation in response to Dabrafenib were included in the model. For each kinase, a score was calculated as the maximum measured log₂ fold change multiplied by the predicted functional score of the phosphorylation site. The final prize per kinase was defined as the maximum score obtained across all its sites. Kinase-substrate links were then aggregated such that literature-derived interactions were assigned a score of 1, while scores from kinase library data and Phosformer predictions were scaled between 0 and 1. When integrating the resources, literature-derived scores were given the highest priority, followed by Phosformer scores, and finally kinase library scores. These values were used as edge weights and further scaled by multiplying with the phosphorylation signal of the respective kinase, restricting the analysis to kinases significantly regulated in the phosphoproteomics time course. To connect kinase-substrate relationships to downstream effects, we constructed an intermediate layer by linking phosphosites to their parent proteins (e.g., MYC_S62 to MYC). This was then complemented with a protein-protein interaction network restricted to transcription factors and TPP-derived proteins. Network reconstruction was performed using a Directed Prize-Collecting Steiner Tree (PCST) approach formulated as an Integer Linear Programming (ILP) problem. The goal was to extract a high-confidence signaling subnetwork that connects the perturbation source to downstream terminal nodes (phosphosites, TFs, TPP hits) while maximizing biological relevance and minimizing network complexity. To enforce rootedness, a virtual "perturbation" node was introduced and connected to the root of the perturbation (BRAF, P15056) with zero cost. The optimization maximizes the trade-off between the collected prizes (measured data relevance) and the cost of the edges included in the solution:

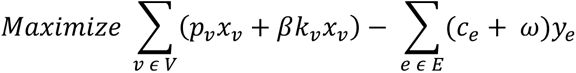

*Where:*

- *x_v_ ∈ {0, 1} and y_e_ ∈ {0, 1} are binary decision variables indicating the selection of node v and edge e, respectively*
- *p_v_ represents the data-derived prize for node v*
- *k_v_ represents the normalized kinase activity score*
- *c_e_ is the intrinsic cost of the edge*
- *β (kinase weight) is a hyperparameter controlling the influence of kinase activity on network selection*
- *ω(cost addition) is a sparsity parameter that penalizes the inclusion of edges to control network size*

The optimization was subject to the following topological constraints:

1. Edge Consistency: An edge can only be selected if both its source and target nodes are selected.
2. Flow Connectivity: Selected receiver nodes (excluding the root) must have at least one incoming edge; selected non-terminal nodes must have at least one outgoing edge.
3. Cycle Elimination: Cycles were prevented using Miller-Tucker-Zemlin (MTZ) type constraints based on node distance from the perturbation source.
4. Fixed Interactions: Known interactions that participate in the canonical BRAF response were constrained to be included in the final solution (BRAF-MAPK1 and BRAF-MAPK3).

The ILP problem was formulated using CORNETO (ref). The mathematical optimization toolkit GUROBI (v11.0) was then employed to solve the problems using a time limit of 30 seconds and mipGap = 0.05.

To compare networks generated from different input sets, we quantified the occurrence of each node across the 63 networks (252 in total). This measure of node importance was then used to weight phosphosites for pathway enrichment analysis at the network level. Node degree was defined as the average total number of edges per node across all networks. To assess the robustness of edge and node inclusion, we applied the columnOverlapCoefficient() function from the ribiosUtils R package, which calculates the pairwise Jaccard index between networks.

